# The human ribosome modulates multidomain protein biogenesis by delaying cotranslational domain docking

**DOI:** 10.1101/2024.09.19.613857

**Authors:** Grant A. Pellowe, Tomas B. Voisin, Laura Karpauskaite, Sarah L. Maslen, Alžběta Roeselová, J. Mark Skehel, Chloe Roustan, Roger George, Andrea Nans, Svend Kjær, Ian A. Taylor, David Balchin

## Abstract

Proteins with multiple domains are intrinsically prone to misfold, yet fold efficiently during their synthesis on the ribosome. This is especially important in eukaryotes, where multidomain proteins predominate. Here, we sought to understand how multidomain protein folding is modulated by the eukaryotic ribosome. We used cryo-electron microscopy and hydrogen/deuterium exchange-mass spectrometry to characterise the structure and dynamics of partially-synthesised intermediates of a model multidomain protein. We find that nascent subdomains fold progressively during synthesis on the human ribosome, templated by interactions across domain interfaces. The conformational ensemble of the nascent chain is tuned by its unstructured C-terminal segments, which keep interfaces between folded domains in dynamic equilibrium until translation termination. This contrasts with the bacterial ribosome, on which domain interfaces form early and remain stable during synthesis. Delayed domain docking may avoid interdomain misfolding to promote the maturation of multidomain proteins in eukaryotes.

## Introduction

Most proteins contain more than one domain^1^. Although functionally advantageous, combining domains into a single polypeptide often compromises refoldability^2–11^, necessitating cellular mechanisms tailored to multidomain protein biogenesis. A fundamental solution is to couple folding to translation on the ribosome. Cotranslational folding shapes protein maturation in several ways. Vectorial synthesis can separate folding into elementary steps^12–14^, while interactions with the ribosome surface can destabilise native folds^15–17^, and tethering to the ribosome can stabilise unique folding intermediates^18–20^. In the case of multidomain proteins, cotranslational folding has been suggested to avoid misfolding by favouring sequential folding of individual domains^21,22^. Nonetheless, domains do not necessarily behave as independent units during cotranslational folding. Detailed studies of the multidomain bacterial protein EF-G have revealed complex interdependencies between folding domains on the ribosome^23–26^. How domain-domain interactions are modulated by the ribosome is poorly understood.

Multidomain proteins are ∼3 times more frequent in eukaryotic compared to prokaryotic proteomes^27^, implying increased evolutionary pressure to optimise their biogenesis. Indeed, multidomain proteins typically fold more efficiently in eukaryotes than bacteria^28^. Experiments in cell-free translation systems have further suggested that eukaryotic and prokaryotic ribosomes differ fundamentally in their ability to promote multidomain protein folding^22,29^. Although they are similar overall, bacterial and eukaryotic ribosomes differ in the architecture of their exit tunnels^30,31^, and several eukaryote-specific ribosomal proteins cluster near the exit port where nascent polypeptides emerge into the cytosol^32^. Whether species-specific features of ribosomes directly influence cotranslational folding is not clear.

To study how eukaryotic ribosomes shape multidomain protein folding, we focused on Firefly Luciferase (FLuc) as a nascent chain model. FLuc is a conformationally labile 2-domain protein, the efficient biogenesis of which strongly depends on the cellular environment. FLuc refolds extremely slowly (t_½_ ∼75 min) from denaturant in vitro, and populates aggregation-prone intermediates^3,33^. Although the Hsp70 chaperone system substantially accelerates FLuc folding (t_½_ ∼4 min)^3,34^, FLuc maturation is optimal when coupled to translation on the eukaryotic ribosome, where its folding is synchronised with synthesis (t_½_ ∼1 min)^21,35^.

Understanding why cotranslational folding is efficient requires molecular insight into nascent chain (NC) conformation. However, the structural dynamics of ribosome-tethered NCs are challenging to resolve^36^, and eukaryotic NCs have not been characterised due to the absence of suitable approaches. Here we extend peptide-resolved hydrogen/deuterium exchange-mass spectrometry (HDX-MS) to human ribosome-nascent chain complexes (RNCs)^19^. In combination with cryo-electron microscopy (cryoEM) and orthogonal biochemical approaches, this allowed us to describe the local conformational landscape and ribosome contacts of NCs at specific points in their synthesis. We find that FLuc (sub)domains fold interdependently on the human ribosome. Folding of the N-terminal domain is templated by its partner interface, promoting native interdomain contacts. Folded domains then detach as translation progresses, potentially helping to avoid entrenching misfolded states. This contrasts with the bacterial ribosome, which permits stable docking of N-terminal domains and interferes with folding of C-terminal domains.

## Results

### Design and preparation of Firefly Luciferase ribosome:nascent chain complexes

FLuc consists of 550 residues across two domains (Fig. 1A)^37^. The large N-terminal domain contains the active site, and is connected via a flexible linker to a smaller C-terminal domain. The N-domain has a complex discontinuous topology, with two inverted Rossman-like folds connected by a β-barrel roll (β-roll). The N-terminal part of the N-domain (Ns, residues 15-190) has been shown to form a protease-stable subdomain during translation but not refolding from denaturant^21^.

**Fig. 1.**
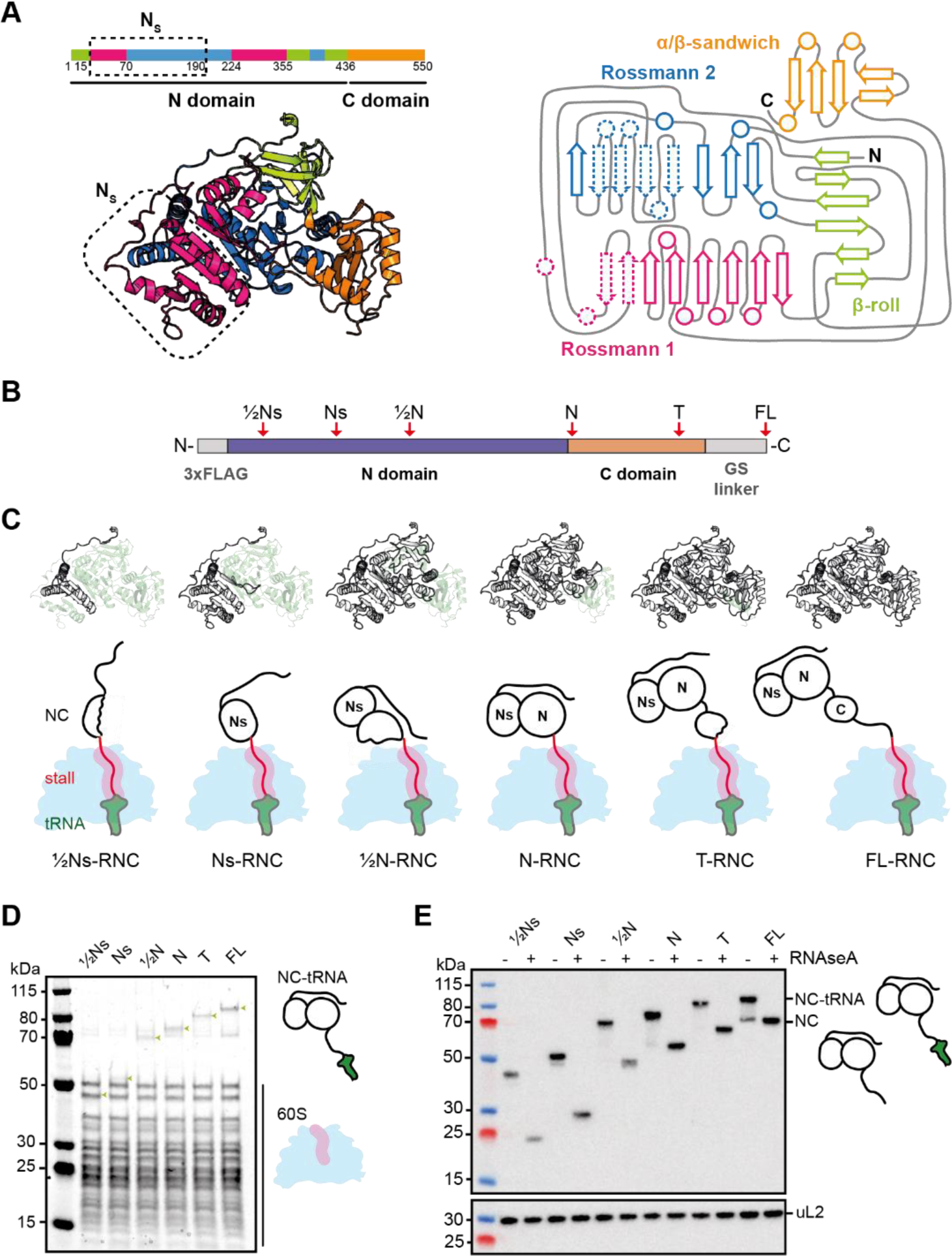
FLuc ribosome:nascent chain complexes. **A,** Structure and domain architecture of FLuc. The structure (left) is predicted by Alphafold 2^60,61^ and the topology map (right) is based on PDB: 1LCI^37^. Elements corresponding to the sub-N domain (Ns) are indicated by dashed lines on the topology map. **B,** Design of FLuc stalling constructs. Positions at which XBP1u+ was inserted are indicated by red arrows. **C**, Schematic diagram of ribosome:nascent chain complexes (RNCs). The fraction of FLuc synthesised in each RNC is coloured white. **D**, Coomassie-stained SDS-PAGE of purified RNCs. Bands corresponding to 60S ribosomal proteins and the NC linked to peptidyl-tRNA (green arrows) are indicated. **E**, Anti-FLuc immunoblot of purified RNCs. RNase/EDTA treatment confirms that the NCs are covalently linked to peptidyl-tRNA. Bottom: immunoblot against ribosomal protein uL2 as a loading control. See also Fig. S1.

To characterise folding intermediates of FLuc on the human ribosome, we first developed an approach to prepare suitable quantities of homogeneously-stalled RNCs, analogous to previous work in bacteria^19,38–40^. To stall translation, we designed an arrest-enhanced variant of XBP1u, guided by a previous mutagenesis study^41^. The resulting XBP1u+ was ∼4-fold more efficient at stalling FLuc translation in human cells than the previously characterised arrest-enhanced variant XBP1u^42^ (Fig. S1A). Stalling positions were chosen based on the domain architecture of FLuc (Fig. 1B). The RNCs exposed half of the Ns subdomain (½Ns-RNC, residues 1-123), the entire Ns (Ns-RNC, 1-208), half of the N-domain (½N-RNC, 1-388), or the entire N-domain (N-RNC, 1-458) (Fig. 1C). To mimic the stage of folding immediately prior to translation termination, we additionally created T-RNC (1-528), in which the C-terminal 22 amino acids (aa) of FLuc was replaced with the stalling sequence. As a control, we designed an RNC where the entire FLuc sequence was extended from the ribosome via a 50 aa GS-rich linker (FL-RNC).

We expressed the stalling constructs in suspension-adapted HEK293 cells and purified RNCs via an N-terminal 3xFLAG tag on the NC. We rigorously purified the RNCs to remove loosely-bound interactors including chaperones, allowing us to isolate the effect of the ribosome on NC folding. High-salt purification also removed the 40S subunit, facilitating downstream HDX-MS analyses, which are limited by sample complexity. NCs resolved by SDS-PAGE were sensitive to RNase, indicating that they were covalently linked to peptidyl-tRNA (Fig 1D,E). FL-RNC was enzymatically active, indicative of native folding (Fig S1B). Mass spectrometry showed that the purified RNCs were substantially depleted of 40S ribosomal proteins, as expected, but retained the NC at near-stoichiometric levels (Fig S1C,D).

Aside from 60S ribosomal proteins and FLuc, the only other abundant proteins in the RNCs were EIF6, which is established to bind isolated 60S subunits^43^, and Hsp70 (HspA1A) (Fig S1E and Table S1). These each copurified at ∼20% occupancy, irrespective of NC length, arguing against NC-specific binding. To locate Hsp70 on RNCs we used crosslinking-mass spectrometry (XL-MS). We identified crosslinks between Hsp70 and the N-terminal part of ribosomal protein eL24, which is distant from the exit tunnel (Fig S2A). eL24 bridges the small and large ribosomal subunits, and its N-terminus is expected to be flexible in the absence of 40S. Crosslinks to eL24 stemmed from the substrate-binding domain and “lid” of Hsp70, consistent with a substrate-like interaction (Fig S2B). Hsp70 did not crosslink to the NC in any sample, suggesting that it is not directly bound to nascent FLuc in the high-salt purified RNCs.

### Structure of Ns-RNC

To further characterise the stalled RNCs, we solved the structure of Ns-RNC to a global resolution of 2.2 Å by cryoEM (Fig 2A, S3A-D and S4A). As observed for rabbit ribosomes stalled on XBP1u^S255A41^, stalling by XBP1u+ does not induce large conformational changes in the 60S ribosome. The P-site tRNA is poorly resolved due to high flexibility in the absence of the small subunit (Fig 2A and S4E). However, we observe clear density for the P-tRNA CCA tail in our consensus map (Fig 2B and S4F), confirming attachment of the NC.

**Fig. 2.**
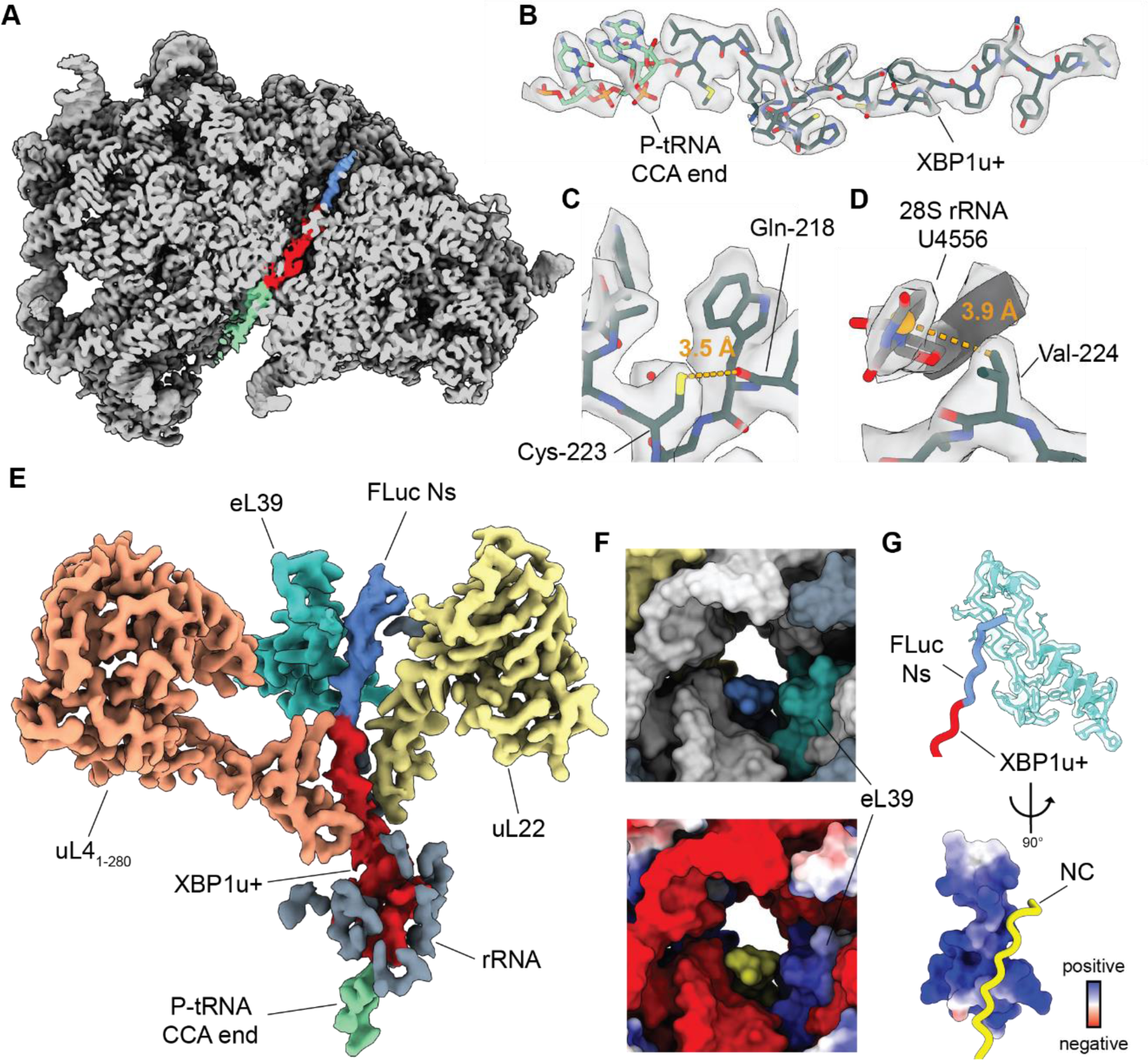
Structure of Ns-RNC. **A,** Cross-section of the consensus map of Ns-RNC, with the 60S ribosome in grey, density corresponding to the P-site tRNA in green, the XBP1u+ arrest peptide in red and FLuc in blue. **B,** Map-model overlay of the XBP1u+ arrest peptide (black) and the CCA end of the P-tRNA (green). **C,** New intra-chain contact in XBP1u+. The thiol sulfur of C223 is within 3.5 Å of the backbone oxygen of Q218 in XBP1u+. **D,** New ribosome contact involving XBP1u+. The centre of the aromatic ring of U4556 (28S rRNA) is within 3.9 Å of the nearest side-chain methyl carbon from XBP1u+ V224. **E,** CryoEM density of the nascent chain within the exit tunnel. XBP1u+ is coloured red and FLuc is in blue. rRNA (grey) is only shown if within 4 Å of the NC. **F,** Top: surface representation of the exit tunnel vestibule showing the NC (blue), uL22 (yellow), eL39 (turquoise), rRNA (white) and other ribosomal proteins (grey). Bottom: same view coloured by electrostatic potential (blue: positive; red: negative) with the NC in yellow. **G,** Top: close-up view of the NC (red: XBP1u+; blue: FLuc) in proximity of eL39. Bottom: view rotated by 90° and coloured according to electrostatic potential (blue: positive; red: negative) with the NC in yellow. See also Fig. S3 and S4, and Table S1.

The stalling sequence is well resolved in our map, allowing us to model all side chains and 2 ordered water molecules near R221 and W226 (Fig 2B an S4B-D). XBP1u+ adopts a similar conformation to that previously observed for XBP1u, including a turn involving residues W219 to W226^41^ (Fig S4H). Compared to wild-type XBP1u, XBP1u+ contains 4 mutations (L216I, Q223C, P224V, S225A; Ns-RNC numbering) that increase its resistance to NC folding-induced release^41^, two of which we can rationalise using our structure. The Q223C mutation places a thiol sulfur within 3.5 Å of the backbone carbonyl oxygen of Q218, allowing the formation of a hydrogen bond that likely stabilises the W219-W226 turn^44,45^ (Fig 2C). The P224V substitution positions one of the side chain methyl carbons within 3.9 Å of the aromatic ring of 28S rRNA residue U4556, compatible with a CH – π interaction^46,47^ (Fig 2D). Thus, stabilisation of the arrest peptide conformation and increased interactions with the ribosome may both contribute to the increased stalling efficiency of XBP1u+.

Our map shows continuous density for the nascent chain in the ribosomal exit tunnel. Clear side chain density allowed us to confidently model the entire XBP1u+ and 8 FLuc residues N-terminal to the stalling sequence (Fig 2B,E and S4G). Despite the volume available to the NC past the constriction point^30,31^, the path of nascent FLuc is biased towards one side of the tunnel, where it occupies a groove formed by rRNA and the eukaryote-specific ribosomal protein eL39 (Fig 2F,G). The positively charged inner surface of eL39 confers a mixed-charge character to the groove, suggesting that the trajectory of the NC may be guided by charge effects (Fig 2F,G). FLuc adopts a partially compacted structure in the exit tunnel, forming a left-handed helix with 3 residues per turn and a ∼3 Å rise per residue, resembling a κ-helix^48^ (Fig 2E and S4G). The same residues make up an unstructured coil in native FLuc, suggesting that confinement in the eL39/rRNA groove may contribute to stabilising non-native secondary structure in the NC.

The absence of clear density for FLuc beyond the exit tunnel indicated that the emerging NC is conformationally dynamic. Consistent with this, we identified frequent crosslinks between NCs and solvent-exposed residues on the ribosome surface (Fig S2C,D). uL29, directly at the tunnel vestibule, crosslinked to all NCs. Longer NCs also crosslinked to ribosomal proteins further from the tunnel exit, including eL22, uL22, uL24 and eL38.

### Sequence of folding events during FLuc synthesis

We next used HDX-MS to probe the conformation of FLuc on the ribosome. We measured peptide-resolved deuterium uptake of NCs, and used isolated (off-ribosome) full-length FLuc (FL-FLuc) as a reference for the native state (Fig 3A). The FLAG tag did not affect deuterium uptake of FL-RNC and was retained (Fig S5A). Sequence coverage of NCs was >83% and most peptides were detected across different RNCs, allowing quantitative comparison between states (Fig S5B and Data S1).

**Fig 3.**
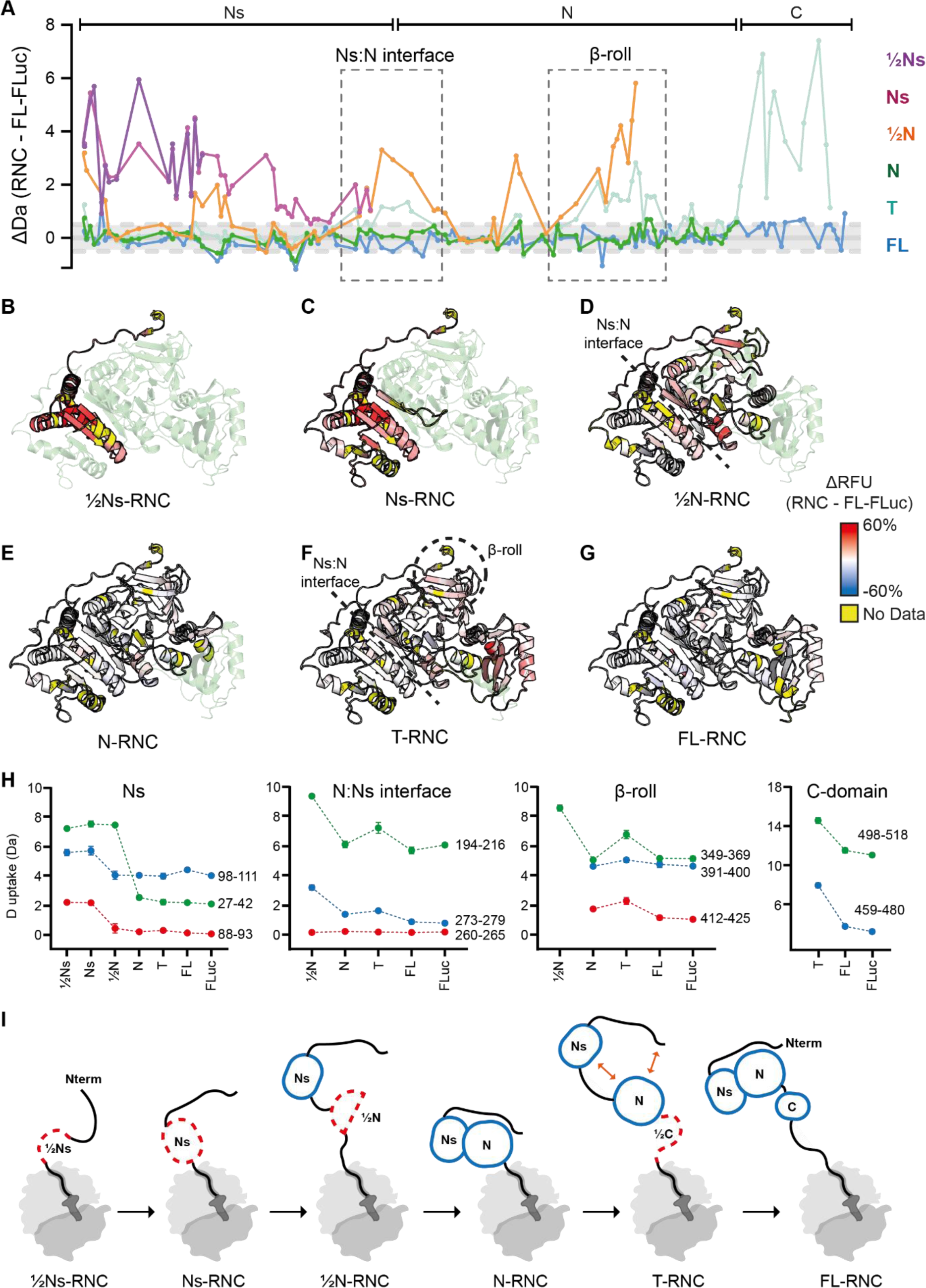
Sequence of folding events during FLuc synthesis. **A**, HDX-MS analysis of FLuc NCs. Difference in deuterium uptake, after 3 min deuteration, between FLuc NCs and native FL-FLuc. Larger values indicate more deuteration of NCs compared to FL-FLuc. Grey dashed lines indicate ±0.5 Da. Peptides forming the Ns:N interface and β-roll are indicated. Data represent the mean of at least 3 independent experiments. **B-G**, Difference in relative fractional uptake (ΔRFU), after 3 min deuteration, between each NC and FL-FLuc. Data are mapped onto the Alphafold2 model FL-FLuc. Darker red indicates increased deuteration of NCs compared to FL-FLuc. Regions without peptide coverage are coloured yellow. **H**, Deuterium uptake after 3 min deuteration, for representative peptides from NC and isolated FL-FLuc (FLuc). Error bars, s.d.; n = 3 independent experiments. Dashed lines are guides for the eye only. **I**, Schematic of FLuc intermediates on the ribosome. Folding of Ns is delayed until the native interface with N is synthesised (½N-RNC). Ns and N then stably associate when N is completely synthesised and close to the ribosome surface (N-RNC). During subsequent C-domain synthesis (T-RNC), Ns:N and β-roll are destabilised. Native interdomain contacts are recovered when the C-domain emerges fully from the ribosome (FL-RNC). See also Fig. S5 and Data S3.

HDX-MS data for each RNC are summarised in Fig 3A-G and representative peptides are shown in Fig 3H. We considered peptides to be natively folded only when they differed in absolute uptake from FL-FLuc by less than 0.5 Da. We found that ½Ns was globally deprotected relative to the same region in FL-FLuc, indicating that it is unfolded (Fig 3B). Ns was also globally unfolded, and peptides from this subdomain reached near-native levels of deuterium exchange only when a larger part of the N domain was synthesised (½N-RNC, Fig 3C,D). The interface between Ns and N remained deprotected until the N-domain was complete (N-RNC), stabilising the β-roll that connects the N domain to the extreme N-terminus of FLuc (Fig 3E). At the final stage of translation, prior to termination, the extreme C-terminal residues of FLuc are within the ribosomal exit tunnel and therefore unavailable to fold with the rest of the NC. In our T-RNC construct which mimics this step, the C domain is highly deprotected relative to FL-FLuc, as expected. Surprisingly, the N domain β-roll and the Ns:N interface of T-RNC are also deprotected compared to RNCs mimicking earlier stages of translation (Fig 3F,H). Extending the C-terminus of FLuc from the ribosome on a 50 aa linker (FL-RNC) resulted in native-like folding of the C-domain, and recovered the stability of the N-domain (Fig 3G). Together, these data resolve length-dependent local folding of partially-synthesised FLuc on the human ribosome (Fig 3I). We find that Ns folding is delayed relative to synthesis, and the NC is destabilised prior to translation termination.

### Folding and docking of the Ns subdomain is delayed relative to synthesis

Our HDX-MS analysis indicated that stable folding of Ns was triggered by partial synthesis of the larger N domain. As an orthogonal approach to identify stable sub-domains, we performed limited proteolysis of RNCs using proteinase K (PK). Ns-RNC was rapidly digested by PK, consistent with our conclusion that it does not fold independently on the ribosome (Fig 4A). Digestion of ½N-RNC resulted in weak accumulation of the previously-described ∼22 kDa intermediate corresponding to Ns^21^, suggesting that Ns is stabilised by interactions with the N domain, and consistent with HDX-MS showing that Ns is near-natively folded at this chain length (Fig 3A,D,H). The Ns intermediate did not accumulate upon digestion of longer RNCs or FL-FLuc. Instead, we observed a stable intermediate consistent in size with the full N domain (Fig 4A). N-RNC was resistant to PK, as expected based on HDX-MS (Fig 3A,E).

**Fig. 4.**
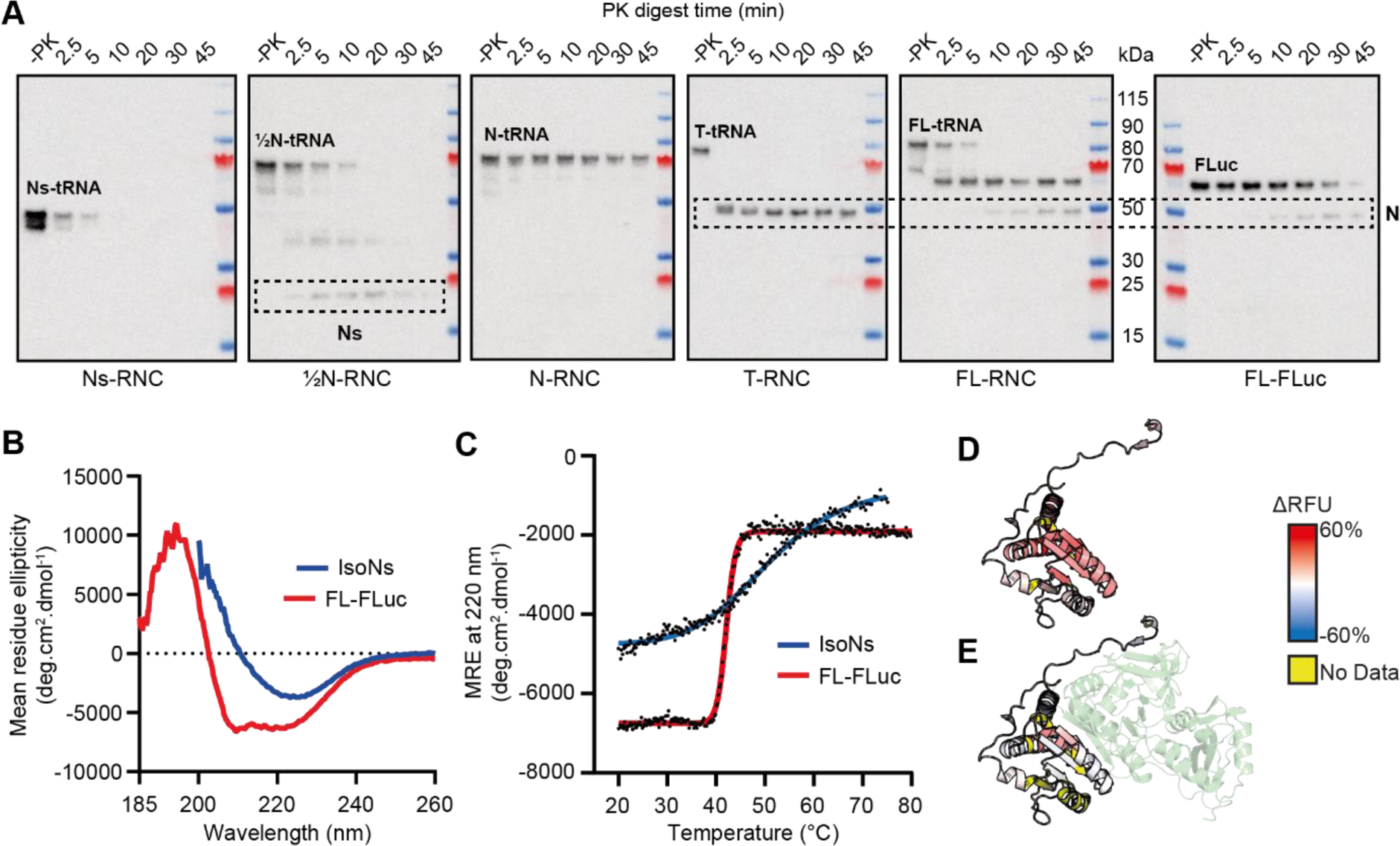
Folding and docking of the Ns subdomain is delayed relative to synthesis. **A,** Limited proteolysis of FLuc RNCs. RNCs were treated with Proteinase K (PK) for different times and probed by immunoblotting using a monoclonal antibody against the FLuc N terminus. **B**, Secondary structure of isolated Ns (isoNs). Circular dichroism spectroscopy of isoNs and FL-FLuc. Spectral deconvolution showed 4.4% α-helix and 38.5% β-sheet for isoNs. Cf. Ns (residues 1-190) in the context of FL-FLuc (PDB: 1LCI): α-helix = 24.7% and β-sheet = 23.5%. **C**, Thermal denaturation of FL-FLuc (T_m_ = 42 °C) and isoNs (T_m_ = 50.5 °C). **D**, Conformational dynamics of isoNs. Difference in relative fractional uptake (ΔRFU), after 3 min deuteration, between isoNs and FL-FLuc. Darker red indicates increased deuteration of isoNs compared to FL-FLuc. **E**, as in D, for Ns-RNC compared to isoNs. Darker red indicates increased deuteration of Ns-RNC compared to isoNs. See also Fig. S6 and Data S3.

To test whether the ribosome modulates Ns folding, we purified isolated Ns (isoNs, residues 1-190) (Fig S6A,B). Unexpectedly, isoNs was dimeric at nM-µM concentrations, as judged by SEC-MALLS and mass photometry (Fig S6C,D). Circular dichroism spectroscopy showed that isoNs had altered secondary structure, with reduced α-helix and increased β-sheet content compared to Ns in the context of FL-FLuc (Fig 4B). Moreover, isoNs showed a shallow transition during thermal denaturation, indicative of low unfolding cooperativity (Fig 4C). Consistent with this, HDX-MS showed that isoNs was globally unfolded relative to native Ns in FL-FLuc (Fig 4D). When compared to Ns-RNC, isoNs was protected at several peptides forming the interface with the N domain in FL-FLuc (Fig 4E). IsoNs may therefore dimerise via the same interface to form a non-native intermolecular β-sheet.

In summary, we find that Ns folds cotranslationally, in agreement with previous reports^21,35^. HDX-MS analysis of purified RNCs and isoNs further showed that Ns is not an independent folding unit, but is held on the ribosome in a folding-competent state until a larger part of the Rossmann fold is available. The Ns:N interface stabilises later, when the entire N domain is synthesised.

### Domain interfaces are destabilised prior to translation termination

Our HDX-MS analysis indicated that the N-domain was natively folded in N-RNC but partially destabilised in T-RNC. As an orthogonal measure of NC stability, we used fluorescein-5-maleimide (F5M) to probe the accessibility of native Cys in FLuc. All Cys are in the N-terminal domain and are buried in the native state (Fig S7A). N-RNC and FL-RNC were labelled by F5M with similar kinetics, whereas T-RNC was labelled significantly faster, consistent with increased Cys exposure (Fig S7B-D). N-domain destabilisation did not depend on the specific C-terminal sequence, as replacing C-domain residues with a Gly/Ser stretch of equivalent length resulted in identical Cys labelling kinetics (Fig S7C,D).

Destabilisation of the N-domain in T-RNC was primarily localised to the β-roll and Ns:N interface (Figure 3A,F,H). Mass spectra for peptides in these regions displayed EX1 kinetics and were multimodal, indicating that both folded and unfolded conformations are populated (Fig 5A,B). In some cases a partially folded population was also evident. Multimodal behaviour was dependent on NC length. Only the unfolded conformation was detected in ½N-RNC, whereas the folded population predominated in N-RNC and FL-RNC. T-RNC showed intermediate behaviour, but was predominantly unfolded. If the unfolded state sampled by T-RNC is in equilibrium with the native state, stabilising the native state would be expected to alter the population distribution. To test this, we added the active site ligands phenobenzothiazine (PBT) and ATP to T-RNC. Note that although T-RNC is not enzymatically active, it has a complete enzyme active site housed in the N domain. Adding ligands protected the β-roll and Ns:N interface regions of T-RNC, FL-RNC and FL-FLuc from deuterium exchange (Fig 5C,D), and shifted multimodal spectra towards the folded population (Figure 5A,B). The high-exchange state(s) populated by T-RNC is therefore in equilibrium with the native state, and not stably misfolded or kinetically trapped.

**Fig. 5.**
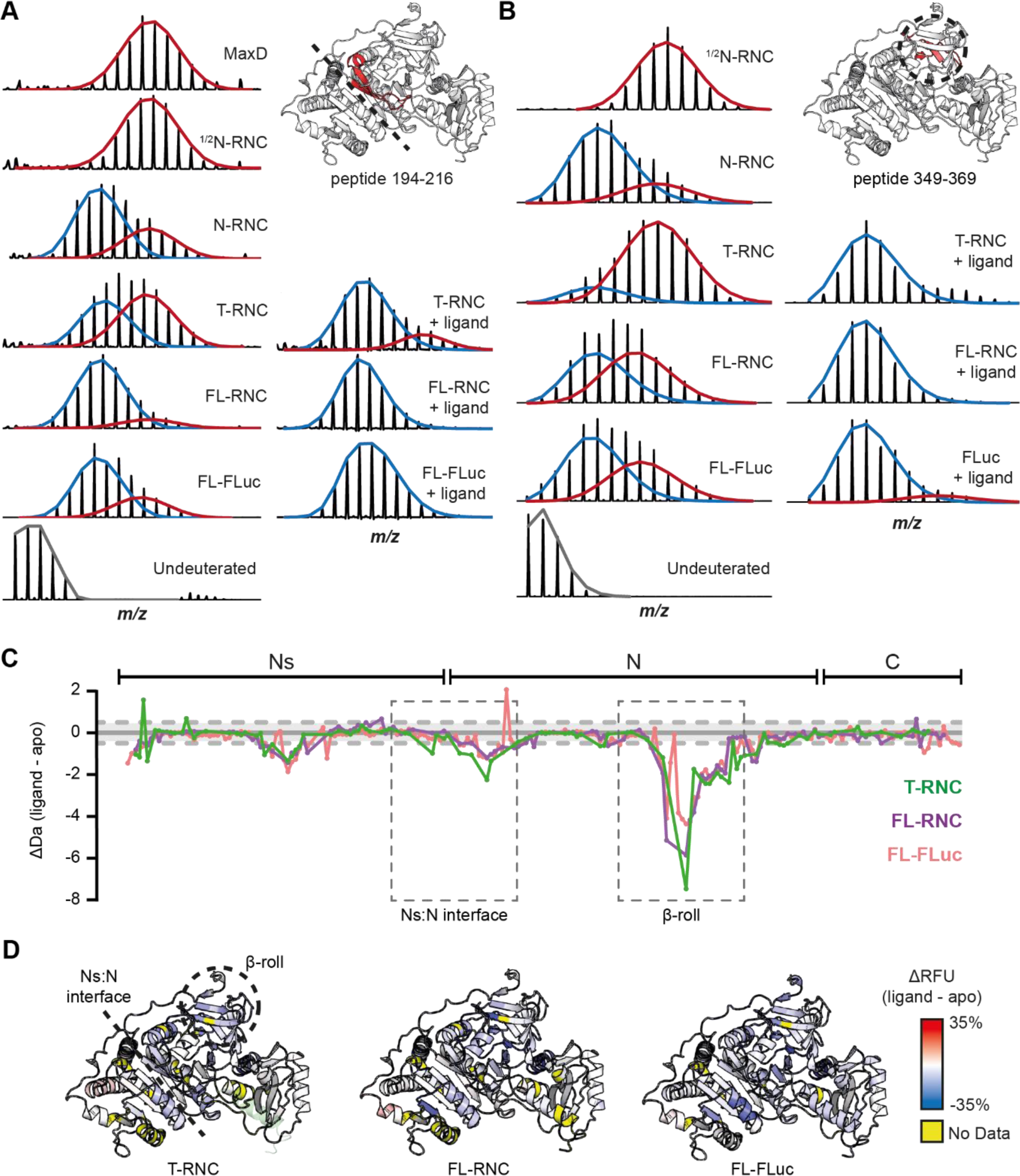
FLuc domain interfaces are destabilised prior to translation termination. Concerted H/D exchange at FLuc domain interfaces. Mass spectral envelopes for peptide 194-216 at the Ns:N interface (**A**) and 349-369 in the β-roll (**B**) of FLuc. Isotopic distributions were fit to single or bimodal gaussian distributions using HX-Express3.0^62^. The high-exchanging population is coloured red and the low-exchanging population is coloured blue. Except for the maximally deuterated sample (MaxD), samples were deuterated for 3 min. In the +ligand condition, samples were incubated with 50 µM PBT and 1 mM ATP prior to deuteration. **C**, Effect of PBT/ATP (ligand) on FLuc conformational dynamics. Difference in deuterium uptake, after 3 min deuteration, between FLuc samples with and without ligand. Smaller values indicate less deuteration of the ligand-bound samples. Grey dashed lines indicate ±0.5 Da. Data represent the mean of at least 3 independent experiments. **D**, Difference in relative fractional uptake (ΔRFU), after 3 min deuteration, between FLuc samples with and without ligand. Regions coloured blue are protected upon ligand binding. See also Fig. S7 and Data S3.

Together, these data show that the conformational ensemble of nascent FLuc is altered by synthesis of a C-terminal unstructured sequence, which shifts the equilibrium towards the unfolded state and destabilises pre-formed domain interfaces.

### Human and bacterial ribosomes differentially affect the NC conformational ensemble

To understand whether cotranslational folding intermediates differ between human and bacterial ribosomes, we generated equivalent bacterial RNCs (bRNCs) for FLuc in *E. coli* (Fig S8A-C). Stalling positions were offset by 12 aa to account for the difference in length of the stalling sequence between XBP1u+ (22 aa) and the bacterial arrest peptide SecM^Str^ (10 aa). HDX-MS showed that the overall sequence of folding events is similar on bacterial and human ribosomes, including delayed folding of Ns (Fig S9A,B). However, N domain peptides were protected on the bacterial compared to human ribosome at all NC lengths, indicating that the environment of backbone hydrogens is altered (Fig 6A,B). Furthermore, the bacterial ribosome modulated the domain docking equilibrium. The Ns:N interface was not destabilised in T-bRNC, indicating that the unfolded conformation is sampled much less frequently than on the human ribosome (Fig 6C and 5A). Similar behaviour was observed for the β-roll (Fig S9C). The bacterial ribosome therefore shifts the conformational equilibrium of the N-domain of nascent FLuc to favour the domain-docked state.

**Fig. 6.**
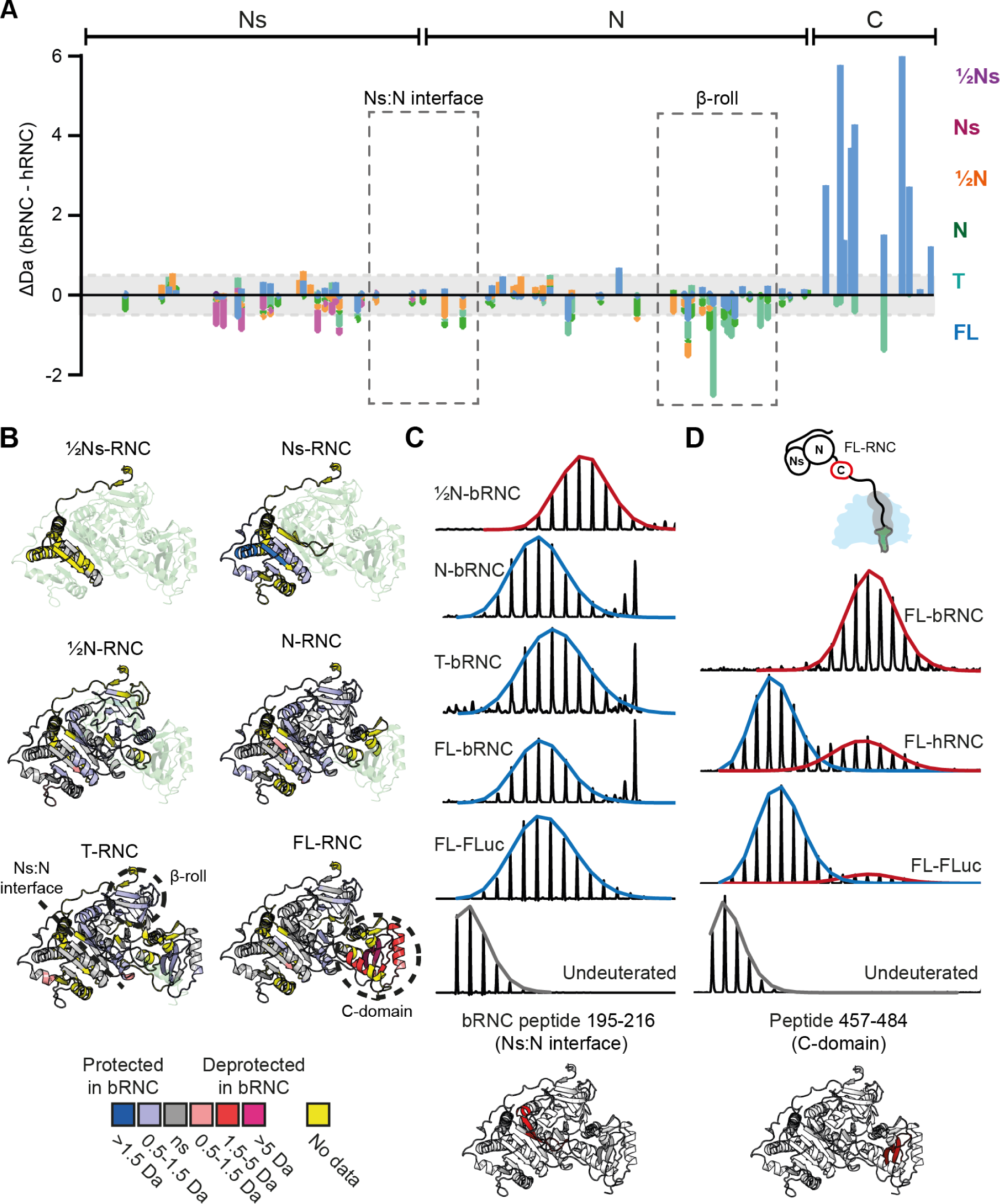
Human and bacterial ribosomes differentially affect the conformational ensemble of FLuc NCs. **A,** Comparison of human and bacterial RNCs. Difference in deuterium uptake, after 3 min deuteration, between FLuc NCs on the human and bacterial ribosome. Positive values indicate more deuteration on the bacterial ribosome. Negative values indicate more deuteration on the human ribosome. **B**, As in A, mapped onto the structure of FL-FLuc. Regions that are protected in bRNCs compared to hRNCs are coloured blue, while deprotected regions are coloured red. Regions without peptide coverage are coloured yellow. **C**, Mass spectral envelopes for peptide 195-216 at the Ns:N interface of bRNCs and FL-FLuc. The high-exchanging population is coloured red and the low-exchanging population is coloured blue. **D**, Mass spectral envelopes for peptide 457-484 in the C-domain of FL-RNCs and FL-FLuc, coloured as in C. See also Fig. S9, Data S3 and Data S5.

The C-domain in FL-RNC showed the opposite effect, and was substantially deprotected on the bacterial compared to human ribosome (Fig 6A,B). Similar to the Ns:N interface and β-roll, peptides in the C domain were bimodal, indicative of a conformational equilibrium between folded and unfolded states (Fig 6D). Whereas the unfolded conformation was rarely sampled on the human ribosome and in isolated FL-FLuc, the same peptide was predominantly unfolded on the bacterial ribosome. The C-domain was nonetheless folding-competent, as FL-bRNC was enzymatically active (Fig S8D). The bacterial ribosome therefore interferes with folding of the C-domain of FLuc near the ribosome surface.

Interactions with the ribosome surface may bias the NC conformational ensemble^15–17,49,50^. To identify interaction sites, we used HDX-MS to analyse the conformational dynamics of human and bacterial ribosomal proteins, in the presence and absence of different NCs (Fig 7A-E and Fig S10A,B). The tunnel-facing surface of eukaryote-specific eL39 on the human ribosome was strongly protected from deuterium exchange, consistent with our cryo-EM structure of Ns-RNC. Protection was not NC-specific, suggesting that the interaction with eL39 does not depend on the sequence of the NC. In addition, NCs protected sites on human ribosomal proteins uL22, eL22, uL29, eL19 and eL31. eL19 and eL31 are unique to eukaryotes and form part of the binding site for the ribosome-associated protein βNAC^51^. In bacteria, the most prominent interaction sites were extended loops on uL23 and uL24. These loops protrude into the vestibule, and are absent from the equivalent human proteins. Together, these data argue that species-specific ribosomal elements bind NCs and potentially modulate their conformation.

**Fig. 7.**
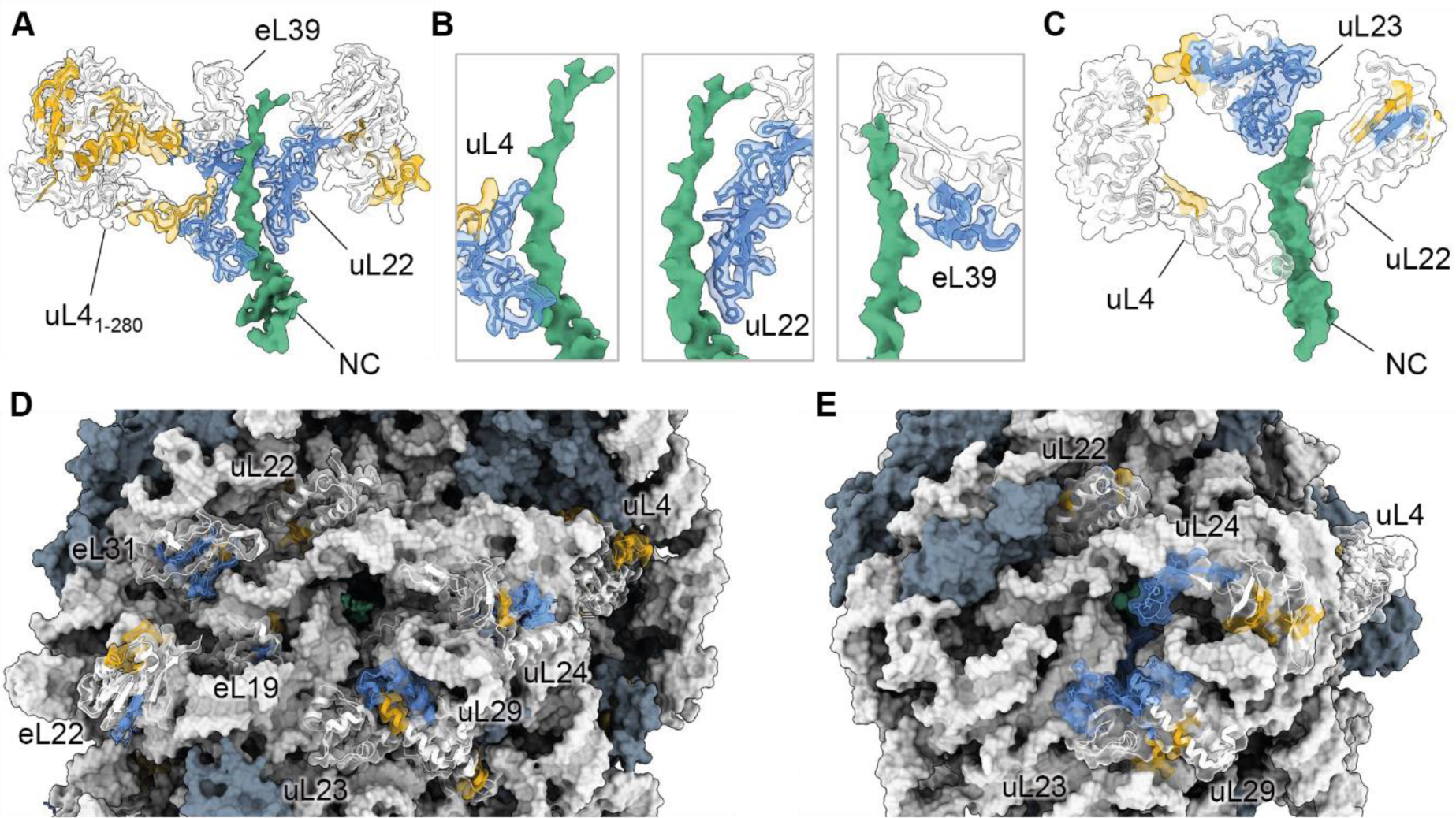
NC interaction sites on human and bacterial ribosomes. **A,** HDX-MS analysis of human ribosomal proteins lining the exit tunnel. Difference in deuterium uptake, after 3 min deuteration, between human RNCs and empty 80S ribosomes. Sites protected by NCs (> 0.5 Da) are coloured blue on the structure of Ns-RNC. Regions that do not change in uptake are coloured white and regions without peptide coverage are coloured yellow. Only residues 1-280 of uL4 are shown for clarity. **B**, Close-up views of uL4, uL22 and eL39, coloured as in A. **C**, HDX-MS analysis of bacterial ribosomal proteins lining the exit tunnel. Difference in deuterium uptake, after 3 min deuteration, between bacterial RNCs and empty 70S ribosomes. Protection is mapped as in A, on the structure of a SecM-stalled ribosome (PDB: 8QOA^63^). **D**, as in A, for proteins on the human ribosome surface. Proteins not analysed are coloured grey. **E**, as in C, for proteins on the bacterial ribosome surface. See also Fig. S10, Data S6 and Data S7.

## Discussion

Here we characterise sequential cotranslational folding intermediates on the human ribosome. We trace the path of the NC through the exit tunnel, identify NC interactions with the ribosome surface, and show how local folding dynamics change during synthesis. This is the first such analysis of eukaryotic RNCs, and reveals differences in NC folding compared to bacterial ribosomes.

We find that FLuc does not fold in a simple domain-by-domain fashion on the ribosome. The Ns subdomain folds only when a larger part of the N-domain is synthesised, then latches onto the N-domain when the β-roll is complete. (Sub)domain folding is therefore stabilised by interactions across domain interfaces. Although Ns initially docks against N, the Ns:N and β-roll interfaces are destabilised during subsequent synthesis of the C-domain. As a result, domain interfaces are maintained in dynamic equilibrium between docked and undocked states, delaying stable domain association until release from the ribosome.

### Optimising multidomain protein biogenesis

Combining domains in a single polypeptide expands protein function but introduces several folding challenges^6,8^. First, domain interfaces must be established with the correct geometry. Second, the co-existence of two unfolded domains entails the risk of interdomain misfolding. These challenges manifest during attempted refolding in vitro, which is complicated by non-native interdomain interactions (Fig. 8A)^3,4,9,52,53^. The same folding vulnerability is illustrated in our study, by the tendency of isoNs to form a non-native dimer when the native interface is unavailable.

**Fig. 8.**
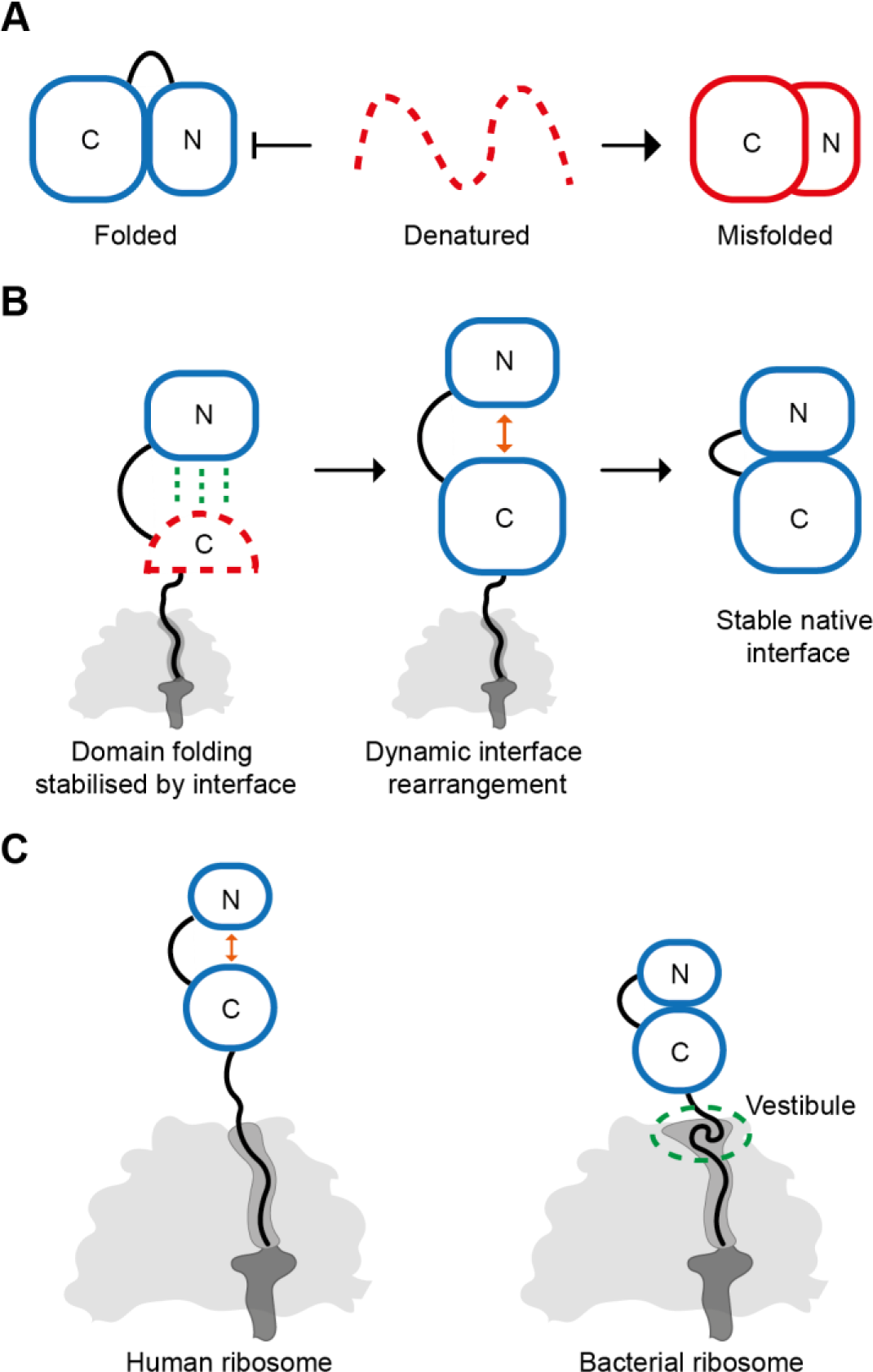
Multidomain protein biogenesis modulated by the ribosome. **A,** Interdomain misfolding in vitro. Denatured multidomain proteins tend to spontaneously misfold in vitro, forming compact non-native states^3^. **B**, Factors promoting multidomain protein biogenesis. Domain folding is stabilised when the partner interface emerges from the ribosome, thereby avoiding misfolding with elements that are not yet synthesised. Later in translation, dynamic rearrangement of domain interfaces on the ribosome may avoid entrenching non-native contacts. **D**, The wide vestibule of bacterial ribosomes accommodates collapsed or partially folded NCs, preventing unstructured C-terminal sequences from destabilising domain interfaces.

How do multidomain proteins fold efficiently in vivo? Previous work has shown that interdomain misfolding can be circumvented by separating domain folding on the ribosome^22^, or rescued by the Hsp70 chaperone system^3,34,54^. Here, we suggest two additional factors that optimise de novo folding of multidomain proteins (Fig. 8B). First, (sub)domain folding is triggered when the native domain interface first becomes available. This might allow the correct interface to form before additional sequence, encoding competing interactions, is synthesised. Second, domain interfaces are partially destabilised by the emergence of unstructured (not-yet-folded) nascent polypeptide from the exit tunnel. A similar phenomenon was observed for EF-G^23^ and DHFR^19^, and may be explained by the entropic cost of tethering a disordered polypeptide to a folded domain^55,56^. This allows for increased sampling of conformational space during synthesis, which could help to avoid kinetic traps or recover from transient non-native interactions between domains. Such a mechanism is in principle sequence-independent (Fig S7C,D), and thus might be a general feature of multidomain protein biogenesis. Chaperones are also involved in FLuc biogenesis^35^, and may play a role in stabilising the nascent Ns before the Ns:N interface is satisfied by native intramolecular contacts. Future research will need to establish how exactly folding on the human ribosome is modulated by molecular chaperones, as well as explore these phenomena for other model proteins.

### Different folding environments on human and bacterial ribosomes

Multidomain proteins often fold more efficiently in eukaryotes compared to bacteria^22,28,29^, but whether the ribosome contributes directly has not been clear. Here, we show how different ribosomes can shape the conformational ensemble of a model NC. In eukaryotes, nascent domains are preferentially undocked and the C-domain folds close to the ribosome. In bacteria, N-domain interfaces are stable throughout translation and the complete C-domain is held in an unfolded conformation. Delayed folding of C-terminal domains may contribute to the general tendency of multidomain proteins to complete folding post-translationally when expressed in bacteria^22,29^.

What differentiates human and bacterial ribosomes? Human ribosome exit tunnels are both narrower and shorter than their bacterial counterparts^30^, likely disfavouring cotranslational folding in the exit tunnel (Fig 7 and 8C). Moreover, although tertiary structure can form in the bacterial exit tunnel as it widens near the exit port^12,57,58^, the equivalent vestibule in humans is constricted by the eukaryote-specific tunnel protein eL39. We show by HDX-MS that diverse sequences directly contact eL39, and by cryo-EM that the NC follows a narrow groove between eL39 and rRNA without sampling substantially compact structures. We speculate that exit tunnel architecture shapes overall NC folding by dictating the conformation of the C-terminal part of the NC. Unstructured C-terminal sequences outside the exit tunnel of the ribosome can destabilise folded domains^19,23,55,56^, thereby amplifying NC interdomain dynamics on the human ribosome (Fig 8C). On the bacterial ribosome, collapse of C-termini in the wide vestibule may insulate already-folded domains from entropic destabilisation (Fig 8C). Indeed, the bacterial ribosome was recently shown to reduce NC entropy to favour partial folding^20^.

Specific interactions with the ribosome surface may also influence folding. On the bacterial ribosome, the NC engages prokaryote-specific loops on uL23 and uL24 that protrude into the vestibule and are established to affect the onset of folding^49,59^. Eukaryote-specific rRNA expansion segments^32^ might also contribute to creating a unique folding environment on the human ribosome.

## Acknowledgements

We thank David Briggs for help with cell culture, Stephane Mouilleron for purified 3C protease, Sergi Garcia-Manyes for the Ulp1 vector, Christelle Soudy for help preparing pepsin beads, Simone Kunzelmann for help with CD deconvolution, Steven Howell for proteomic analysis, Geoff Kelly for preliminary NMR analysis, Emma Couves for cryoEM advice, and all members of the Protein Biogenesis Lab for help and discussion. This work was supported by funding from the UKRI (FoldingMap, EP/X020428/1), and the Francis Crick Institute which receives its core funding from Cancer Research UK (CC2025, CC1063, CC1068), the UK Medical Research Council (CC2025, CC1063, CC1068), and the Wellcome Trust (CC2025, CC1063, CC1068).

## Author contributions

G.P. performed the biochemical and HDX-MS experiments and analysed the data. T.V. collected and analysed the cryoEM data. L.K. contributed to optimising RNC purification. A.R. contributed to optimising HDX-MS workflows. C.R. and S.K. assisted with cell culture, R.G. performed mass photometry, S.L.M. and J.M.S. collected and processed the XL-MS data. A.N. performed Krios data collection. I.A.T performed SEC-MALLS. D.B. conceived and supervised the project, and wrote the manuscript together with G.P. and T.V.

## Supplementary information

**Table S1.** Cryo-EM data collection, map reconstruction and model refinement

**Data S1.** MS analysis of hRNC composition

**Data S2.** XL-MS

**Data S3.** HDX-MS analysis of hRNCs

**Data S4.** MS analysis of bRNC composition

**Data S5.** HDX-MS analysis of bRNCs

**Data S6.** HDX-MS analysis of human ribosomal proteins

**Data S7.** HDX-MS analysis of bacterial ribosomal proteins

## Methods

### Plasmids and constructs

Stalling constructs were synthesized in mammalian expression vector pTwist CMV BG WPRE Neo (Twist Bioscience). The constructs contained an N-terminal 3xFLAG tag and 3C protease site upstream of Firefly Luciferase (subcloned from pGL2^+^), and a downstream stalling sequence XBP1u+ (VPYQPPFICQWGRHCVAWKPLMN). Compared to XBP1u^42^, XBP1u+ contains the following mutations: L216I, Q223C, P224V, and S225A (Ns-RNC numbering). The ORF is under transcriptional control of the CMV promoter, and beta-globin and Woodchuck Hepatitis Virus (WHV) post-transcriptional regulatory elements were included to enhance protein expression. pTwist vectors were amplified in DH5α strain *Escherichia coli* in Luria-Bertani (LB) media and maintained with 100 μg.mL^-1^ ampicillin. Truncations and modifications of the stalling construct were produced by deletion PCR using a Q5-SDM kit or Gibson assembly (New England Biolabs). DNA for transfection was prepared using PureLink™ HiPure Plasmid Maxi-prep kit (ThermoFisher) from 250 mL cultures.

### Human RNC purification

#### Transient transfection and cell harvest

Expi293F suspension cells (ThermoFisher) were maintained at 0.4x10^6^ cells.mL^-1^ twice a week in FreeStyle™ 293 Expression Medium (ThermoFisher) at 37 °C, 8% CO_2_ and shaking at 125 rpm in vented Erlenmeyer flasks (Corning). Maximum volumes for culture were as follows: 30 mL culture in 125 mL flask, 50 mL in 250 mL flask, 100 mL in 500 mL flask, 200 mL in 1 L flask, 600 mL in 2 L flask. Cell number and viability were estimated using a ViCell-XR cell counter (Beckman Coulter) and transfections were only carried out if viability was >95%. Cells were discarded at passage 20. Cells were split to 2x10^6^ cells.mL^-1^ one day prior to transfection and on the day of transfection were diluted to 3.3x10^6^ cells.mL^-1^ in 9/10ths the final transfection volume (3x10^6^ cells.mL^-1^ final cell count). Polyethylenimine (PEI)-DNA complexes were mixed at 3:1 mass ratio, with 1 μg plasmid DNA per 1 mL of 3x10^6^ cells.mL^-1^ cells to be transfected. PEI (Polysciences) (1mg/mL stock, pH 8.0) was diluted to 30 μg.mL^-1^ in half of 1/10th final transfection volume into Opti-MEM Reduced Serum Medium with Glutamax (ThermoFisher), and plasmid DNA was diluted into the same volume of OptiMEM, and were incubated for no longer than 5 min at room temperature (RT). The diluted PEI and DNA were mixed and incubated for 20 min at RT and then added dropwise with swirling to the transfection ready cells and incubated as above for expression of RNCs. Cells were harvested after 24 h of transient expression by centrifugation at 300g. Cells were subsequently washed in ice-cold PBS with cOmplete EDTA-free protease inhibitor tablet (Roche). Harvested cells were used immediately for RNC purification.

#### Cell lysis and affinity purification

All buffers were made using RNase/DNase-free molecular biology grade H_2_O.

Cells were weighed and gently resuspended at 1/100th cell culture volume in ice-cold 80S lysis buffer (50 mM Tris-HCl, pH 7.5, 0.5% NP40-alternative, 5 mM MgCl_2_, 25 mM KCl, 0.2 M arginine-HCl, 1 mM phenylmethylsulfonyl fluoride (PMSF), 1 mM Dithiothreitol (DTT), 2 μL/10 mL RNasIN and EDTA-free cOmplete protease inhibitor tablet) with a Pasteur pipette. RNase-free DNAseI (Qiagen) was added at 10 μL/1 g wet-cell weight and cells were incubated on ice for 30 min to promote lysis. Cell debris was removed by centrifugation at 14,000g for 30 min at 4 °C and the clarified lysate was layered onto a 35% sucrose cushion (35% w/v RNA/DNAse-free sucrose, 50 mM HEPES-NaOH pH 7.5, 500 mM KCl, 5 mM MgCl_2_, 0.2 M arginine-HCl, 1 mM PMSF, and RNaseIN) to enrich for ribosomes. Layered samples were centrifuged at 70,000 rpm in a Beckmann TLA-110 rotor for 2 h, or 55,000 rpm for 4 h in Beckmann Ti-70 rotor at 4 °C. Supernatants were discarded and the clear, straw-coloured pellets containing ribosomes and RNCs were resuspended with gentle agitation overnight in RNC buffer (20 mM HEPES-NaOH pH 7.5, 500 mM KCl, 5 mM Mg(OAc)_2_, 10 mM NH_4_Cl, 10% v/v glycerol, 1 mM PMSF and 0.2 M arginine-HCl) at 4 °C.

Resuspended ribosomal pellets were incubated with RNC buffer-equilibrated agarose ANTI-FLAG® M2 Affinity Gel (ThermoFisher) for 3 h at 4 °C on a spinning wheel (50 μL beads for 50 mL culture). The unbound fraction was collected with a 7,000g centrifugation step at 4 °C along with subsequent washes (3x 15 min) of RNC buffer. 3xFLAG-tagged RNCs were eluted from the resin with 1x 150 ng.μl^-1^ 3xFLAG peptide (Peptide Chemistry STP, Francis Crick Institute) diluted in RNC buffer for 1 h, 4 °C.

#### Generation and purification of 70S RNCs

bRNCs were prepared using construct design and high salt purification methods detailed previously^19,65^ with some minor changes. *Δtig* BL21 (John Christodoulou, University College London) were transformed with RNC constructs containing arrest-enhanced SecM stalling sequence (WWWPRIRGPPGS)^66^, and were grown in ZYM-5052 autoinduction media at 37 °C for 4 h, before cooling to 18 °C for expression overnight. bRNCs were truncated at the following positions: ½Ns-bRNC: 1-135, Ns-bRNC: 1-220, ½N-bRNC: 1-400, N-bRNC: 1-470, T-RNC: 1-540, FL-bRNC: 1-550+GS_50_-SecM. FLuc length was altered to account for the relative lengths of SecM and XBP1u+ in order to maintain total construct length between orthogonal RNCs. All purification buffers additionally contained 0.2 M Arginine-HCl and were bound to, and eluted from ANTI-FLAG® M2 Affinity Gel, as per human RNC purifications.

### Purification of 80S ribosomes

Crude 80S ribosomes were purified as human RNCs with some minor differences. Lysis was performed in 0.5 M KCl lysis buffer without arginine. Additionally, 2.5 mM puromycin was added during lysis to remove nascent chains from ribosome. Cell debris was removed by centrifugation at 14,000g for 20 min, then ribosomes were isolated from the supernatant by sucrose cushion ultracentrifugation as described above. Ribosome pellets were washed twice, and resuspended in RNC buffer.

### RNC quality control

RNC concentration was estimated using A_260_, where 1 A_260_ = 20 pmol/mL = 0.02 μM of 80S ribosomes^67^, or 1 A_260_ = 24 pmol/mL = 0.024 μM of 60S or 70S ribosomes^38^. The mass of 80S ribosomes was calculated using a molecular weight of 4.5 MDa. RNCs were aliquoted and flash frozen for storage at -80 °C.

RNC integrity was checked by running 1-2.5 pmol of RNC in reducing Laemmli Sample Buffer (BioRad) on 4-12% Bis-Tris gels (ThermoFisher) in MES-SDS running buffer. Samples were not boiled. 0.5 μL of 1/10 diluted RNaseA and 0.5 μL of 0.5 M EDTA pH 8.0 was added to a 10 μL RNC sample to digest tRNA, for confirmation of ribosome stalling^38^. Gels were either stained with Coomassie to check characteristic 60S or 80S ribosomal banding, or transferred (BioRad trans-blot turbo) to pre-packed nitrocellulose membranes (BioRad) for Western blot. Nascent chains were blotted using anti-N-terminal FLuc (Abcam, ab185923, lot# GR3317915-2). Membranes were blocked at RT for 1 h, or overnight at 4 °C with 1% (w/v) milk in PBS with 0.1% (v/v) Tween20 (PBST). Antibodies were used at 1:1,000-10,000 dilution, with a Goat anti-Rabbit HRP secondary antibody (Abcam, ab205718) at 1:10,000 after first washing excess primary away with PBST. Membranes were exposed using SuperSignal™ West Pico PLUS Chemiluminescent Substrate (ThermoFisher) on an Amersham 600 imager.

### Purification of recombinant full-length firefly luciferase (FL-FLuc) and off-ribosome truncations

His-SUMO-FLuc was cloned into a pET19 vector for expression in *E. coli.* Isolated truncations were produced by deletion PCR using Q5 site-directed mutagenesis kit (New England Biolabs) and contained an N-terminal His-SUMO tag. His-SUMO-FLuc constructs were transformed into BL21 cells and a single colony was used to seed a starter culture of LB maintained with 100 μg.mL^-1^ ampicillin. The starter culture was used to inoculate ZYM-5052 autoinduction media^68^ with 100 μg.mL^-1^ ampicillin, which was incubated for 4-6 hrs (until clouded) at 37 °C before cooling to 18 °C for expression over 12-16 h. Cell pellets were harvested and washed in PBS before resuspension in lysis buffer (10 mM Tris pH 8.0, 10 mM imidazole, 300 mM NaCl, 5 mM β-mercaptoethanol, 1 mM PMSF, and EDTA-free cOmplete protease inhibitor tablet) and storage at -80 °C.

For purification using an AKTA Pure™ system (Cytiva), resuspended pellets were thawed and incubated for 20 min at RT with 5 μL of Benzonase nuclease (Sigma). Cells were lysed with one pass through a cell disruptor (Constant Systems) at 25 kPsi at 4 °C, and cell debris was removed with a 66,000g centrifugation for 30 min, 4 °C. Supernatants were loaded onto a 5 mL HisTrap HP column (Cytiva) equilibrated with binding buffer (10 mM Tris, pH 8.0, 10 mM Imidazole, 300 mM NaCl, 5 mM β-mercaptoethanol, 1 mM PMSF). Non-specifically bound proteins were removed by washing with 10 mM imidazole in binding buffer, before elution over a 25 CV gradient from 10 to 300 mM imidazole. Fractions containing FLuc were treated with 8 μg Ulp1 protease (prepared in-house) per 2 mg of protein during dialysis overnight into gel filtration buffer (20 mM Tris pH 7.5, 150 mM NaCl, 5 mM β-mercaptoethanol and 0.2 M Arginine-HCl). Cleaved His-SUMO was removed using reverse IMAC in binding buffer containing no imidazole. Eluted fractions containing pure FLuc were concentrated using Vivaspin 50 kDa protein concentrators (Sigma) and injected directly onto a 16/60 Superdex 200 pg column (Cytiva) equilibrated in gel-filtration buffer for further clean-up. FLuc fractions of >95% purity were pooled and buffer exchanged into FLuc storage buffer (25 mM Tris-acetate pH 7.8, 1 mM EDTA, 0.2 M ammonium sulphate, 15% glycerol, 30% ethylene glycol, 2 mM TCEP) and flash frozen for storage at -80 °C.

Before use, the protein was buffer exchanged into the appropriate assay buffers using 7K Zeba spin desalting columns (ThermoFisher) as per manufacturers instruction, and the product centrifuged at 21,000g for 20 mins, 4 °C to remove potential aggregated protein.

### Luciferase activity assays

RNC or FLuc were diluted to 0.2 μM in 10 μL reactions with assay buffer (20 mM HEPES-pH 7.5, 100 mM KCl, 5 mM Mg(OAc)_2_, 1 mM DTT and 0.2 M Arginine-HCl) and equilibrated at 25 °C for 10 mins with either 5 % (v/v) DMSO, 50 μM 2-phenylbenzothiazole (PBT) in DMSO, or 12.5 μg/mL RNaseA. 50 μL luciferase assay reagent (Promega) was added to each reaction and luminescence was immediately measured using a Glomax 20/20 luminometer (Promega) with delay time of 2 s. The relative luminescence units (RLU) were corrected for pmol of protein in each reaction, calculated using A_260_ (for RNC) or A_280_ (for FLuc) measurements. Post-reaction samples were concentration-normalised for gel loading, and immunoblotted for FLuc nascent chains (anti-N-terminal FLuc) to confirm tRNA-bound nascent chains remained intact.

### ProteinaseK digestion assays

RNC or FLuc at 0.1 μM in HDX labelling buffer was treated with 0.5 μL of 8 mg.mL^-1^ 3C protease for 30 min on ice to remove 3x FLAG. 4 μL of 5 mM PMSF was pre-chilled on ice to quench ProteinaseK (PK, Sigma). 30 μL of cleaved RNCs were incubated at 10 °C, 4 μL of the reaction was removed and quenched into 4 uL PMSF as a -PK control. 3 μL of PK was added to the reaction at a concentration of 1 ng.μL^-1^ to initiate the digest reaction. At 2.5, 5 10, 20 and 30 mins, 4 μL was removed from the reaction and quenched in pre-chilled PMSF. 8 μL of reducing Laemmli loading dye was added to each sample, and proteins were separated on 4-12% Bis-Tris SDS-PAGE gels, transferred to nitrocellulose membranes, and Western blotted for FLuc. Amounts of tRNA-bound FLuc remaining at each timepoint was measured by densitometry using ImageJ^69^, and normalised to the -PK control. Data was plotted as a single-phase decay using Graphpad Prism 10.

### Cysteine labelling

RNC or FLuc were buffer exchanged using Zeba spin 7K desalting columns (ThermoFisher), or diluted to 0.1 μM into HDX labelling buffer containing no DTT. For urea denatured controls, 8M urea was included in the HDX labelling buffer, and incubated at RT for 1 h to unfold the protein. Stocks of Fluroesceine-5-malemide (F5M) (ThermoFisher) were produced in dimethylformamide, and diluted 200-fold into HDX buffer prior to equilibration at 10 °C. Labelling was initiated by mixing 3 μL of F5M (0.1 mM final) with 30 μL of sample, and incubated at 10 °C in a Thermomixer (Eppendorf) with gentle agitation at 300 rpm. After 30 s, 2.5 min, 10 min, 30 min and 60 min, 5 μL of the reaction was quenched with 5 μL Laemmli sample buffer containing β-mercaptoethanol. Samples were resolved on a 15-well 4-12% Bis-Tris SDS-PAGE gel in MES-SDS running buffer. Fluorescently labelled bands were detected on a Typhoon-9500 with 488 nm laser excitation, and 750 gain. No background correction was applied.

Labelled tRNA-bands were identified by Western blot of the same gel when probed for luciferase. Any loading errors were identified here and omitted from further analysis. Raw counts from fluorescent gels were quantified using ImageJ^69^, normalised to largest degree of labelling in each reaction, and fitted to a 1-phase exponential plateau in Graphpad Prism 10. Labelling rates were calculated for individual reactions, and plotted for comparison between samples.

Statistical significance between labelling rates was tested using an ordinary 1-way ANOVA between different conditions from at least three independent biological repeats. All data presentation and statistical analysis was carried out in Graphpad Prism 10.

### Cryo-electron microscopy

#### Grid preparation and data collection

Lacey carbon Au 300-mesh grids (Agar Scientific) were glow-discharged in air for 30 s with a GloQube instrument (Quorum Technologies). 6 µl of purified Ns-RNC at a concentration of 0.1 µM were applied to the grids before blotting with a Vitrobot Mark IV (ThermoFisher Scientific) at 22.5 °C and 80% with a blot time of 3 s and a blot force of –1. Grids were plunge-frozen in liquid ethane and stored in liquid nitrogen.

Movies were collected on a 300 kV Titan Krios electron microscope (ThermoFisher Scientific) with a Falcon 4i direct electron detector (ThermoFisher Scientific) and a Selectris energy filter (ThermoFisher Scientific) operating at a slit width of 10 eV. 35,616 movies were collected with EPU v3.8.1.7603 using aberration-free image shift and fringe-free illumination at a nominal magnification of 130,000× for a pixel size of 0.95 Å/px. Data collection details are in Supplementary Table S1.

#### CryoEM image processing

CryoEM image processing is summarised in Fig. S3. Movies were motion-corrected in Relion 4^70^ using MOTIONCOR2^71^ and CTF parameters were estimated with CTFFIND4^72^. Exposures were then transferred to Cryosparc^73^ v4.4.1 and curated based on CTF fit and overall motion. 29,080 exposures were selected and particle picking was performed in Cryosparc with the Blob picker utility. Particles were downsampled by a factor of 4 and extracted before two rounds of 2D classification. Particles from well-resolved 2D classes were selected for ab initio reconstruction (one class) and homogeneous refinement. Particles were re-extracted with a downsampling factor of 2 and used for homogeneous refinement followed by heterogeneous refinement (four classes). Particles refining into a class with obvious signs of alignment on the Lacey carbon edge were excluded from further processing while good particles were retained for homogeneous refinement followed by 3D classification (6 classes, 2 O-EM epochs, 6 Å target resolution, PCA initialisation). Particles from the two best-resolved classes (788,554 particles) were taken forward for processing in Relion 5, where they were extracted with a downsampling factor of 2 and subjected to 3D refinement with Blush regularisation^74^. This map was used as a reference to visualise flexibility of the P-site tRNA by performing 3D classification (4 classes, T=100, 10 iterations, Blush regularisation, no alignment) using a mask around the tRNA binding site (Fig. S4). All particles from the reference map were re-extracted without downsampling and subjected to 3D refinement with blush regularisation. Post-processing, CTF refinement and particle polishing were carried out before 3D refinement (with Blush regularisation) to yield the final consensus reconstruction.

#### Model building

The model of the human 80S ribosome (PDB 6QZP^75^) was rigid-body fitted into the consensus map in WinCoot^76^ v0.9.8.93. Regions of the model corresponding to the small subunit or regions that were absent in our map were removed and the XBP1u model and CCA tail of the P-site tRNA (PDB 6R5Q^41^) were rigid-body fitted independently into the density. The relevant XBP1u+ mutations (L216I, Q223C and P224V) were introduced and the eight C-terminal residues of FLuc were added in WinCoot. Two ordered water molecules were modelled into clear density near the nascent chain (interacting with the backbone nitrogen of R221 and carbonyl oxygen of W226, Fig. S4F-G). Density for a third water molecule was seen near the side-chain of K227 but the map quality was not high enough for unambiguous assignment of the water and side chain densities (Fig. S4H) so the water molecule was not modelled. The entire model was locally refined using Isolde^77^ v1.7.1 in ChimeraX^78^ v1.7.1. Ions and water molecules were modelled and checked in WinCoot manually. The model was subjected to real-space refinement with phenix.real_space_refine (Phenix^79^ v1.21) for eight macro cycles using default parameters, with a custom covalent bond between the carbonyl carbon of XBP1u+ M230 and the 3’ oxygen of P-tRNA A76. . Model validation was performed in Phenix v1.21 (Table S1).

### Circular dichroism spectroscopy

Isolated Ns and FLuc were buffer exchanged into 25 mM Na-phosphate, pH 7.8, 10 mM Mg(OAc)_2_ and 2 mM DTT using 7k Zeba spin columns. Isolated Ns was diluted to 0.15 mg.mL^-1^, and scanned in a 1 mm quartz cell from 260 nm to 200 nm in 0.2 nm increments using a Jasco J-815 CD spectrometer at 20 °C. FLuc at 0.82 mg.mL^-1^ was scanned in a 0.2 mm demountable cell from 260 nm to 180 nm in 0.2 nm increments. For both proteins, 25 scans were taken and averaged, and corrected using a buffer blank recorded in the same cell. Machine units were converted to mean residue ellipticity (MRE) using an average mean residue weight (MRW) of 110 g.mol^-1^. Spectra were deconvoluted using the CDSSTR programme^80^, and compared against the SPD40 protein database. Secondary structure content was estimated from the PDB: 1LCI for FLuc^37^ and isolated NS in the context of folded FLuc using the KCD server^81^.

For thermal melts, both proteins were diluted to 0.15 mg.mL^-1^ in a 1 mm quartz cell. CD was measured at 220 nm over a temperature ramp from 20 °C to 75 °C for Ns, and 20 to 95 °C for FLuc.

### SEC-MALLS

Size exclusion chromatography coupled to multi-angle laser light scattering (SEC-MALLS) was used to determine the oligomeric state of isolated Ns. 100 μL of Ns at 0.1 or 0.2 mg.mL^-1^ was applied to a Superdex 200, 10/30 increase GL column (Cytiva) equilibrated in 20 mM HEPES, pH 7.5, 100 mM KCl, 5 mM Mg(OAc)s, 200 mM Arginine.HCl, 1 mM DTT and 2 mM Na-Azide at a flow rate of 1 mL.min^-1^. Scattered light intensity and protein concentration were recorded using a DAWN-HELIOS-II laser photometer and an OPTILAB-TrEX differential refractometer respectively. The weight-averaged molecular mass of the eluate was determined using combined data from both detectors in the ASTRA software.

### Mass photometry

Isolated Ns was diluted to 50 nM in HDX labelling buffer (20 mM HEPES-pH 7.5, 100 mM KCl, 5 mM Mg(OAc)2, 200 mM Arginine.HCl and 1 mM DTT). Solution phase masses were measured on a TwoMP mass photometer (Refeyn). Videos were recorded on pre-cleaned, high sensitivity microscope slides. Mass calibration was determined using urease, BSA and aldolase covering a mass range of 66 to 544 kDa. Each standard was used at 2-5 nM and diluted 10-fold into PBS for data acquisition. For Isolated Ns, 50 nM protein was diluted 10-fold in the HDX labelling buffer. Videos were recorded for 1 min, and analysed using DiscoverMP v2.5 software.

### Crosslinking-mass spectrometry

RNCs (approximately 200 μL of 0.4 μM) for XL-MS were buffer exchanged into XL buffer (20 mM HEPES, pH 7.5, 100 mM KCl, 5 mM Mg(OAc)_2_ and 1 mM DTT) using Zeba 7K desalting columns. For ligand bound conformations, 50 μM PBT and 5 mM ATP were included in this buffer. Crosslinking was initiated by the addition of 1 mM MS-cleavable disuccinimidyl dibutyric urea (DSBU) (ThermoFisher) at 25 °C for 1 h with gentle agitation. Crosslinking was quenched with the addition of 20 mM Tris-HCl, pH 8.0 and incubated for a further 15 min. XL-MS data was collected and processed as described previously^65^. Crosslinked proteins were reduced (10 mM DTT), alkylated (50 mM iodoacetamide) and digested with Trypsin (ratio of 1:100 enzyme:substrate, 1 h 25 °C) followed by a further digestion (ratio of 1:20 enzyme:substrate, 18 h, 37 °C). Tryptic peptides were fractionated into 5 fractions (10%, 20%, 30%, 40% and 80% acetonitrile in 10 mM NH_4_HCO_3_, pH 8) using high pH reverse phase chromatography on TARGA C18 columns (Nest Group Inc), lyophilized, and resuspended in 1 % formic acid and 2 % acetonitrile. Samples were then subjected to nano-scale capillary LC-MS/MS analysis using the Vanquish Neo UPLC (ThermoScientific Dionex) at a flow rate of 250 nL/min, a C18 PepMap Neo nanoViper trapping column (5 μm, 300 μm × 5 mm, ThermoScientific Dionex), an EASY-Spray column (50 cm × 75 μm ID, PepMap C18, 2 μm particles, 100 Å pore size, ThermoScientific), and quadrupole Orbitrap mass spectrometer (Orbitrap Exploris 480, ThermoScientific). Acquisition parameters were set to data-dependent mode using a top 10 method, recording a high-resolution full scan (R=60,000, m/z 380-1,800) followed by higher energy collision dissociation of the 10 most intense MS peaks, excluding ions with precursor charge state of 1^+^ and 2^+^.

For data analysis, Xcalibur raw files were converted to MGF format using proteome discoverer 2.3 and used directly as input files for MeroX ^82^. Searches were performed against an *ad hoc* protein database containing sequences of the 80S ribosome, HSPA1A, EIF6 and FL-RNC, as well as a set of random decoys sequences generated by the software. Crosslinks were analysed using XiView^83^, and intra-FLuc XLs visualised using PyLink Viewer^84^. Crosslinks were filtered by match score > 50.

### Mass spectrometry analysis of RNC composition

RNCs or ribosomes were purified as above in high salt RNC buffer and processed and collected as detailed previously^19,65^. 5-10 μg of RNC or protein was separated in 8 mm on NuPAGE 1mm 12% Bis-Tris 10 or 12 well SDS-PAGE gels prior to Coomassie Blue staining (Quick Coomassie Stain, Generon). Bands were destained in extraction buffer (50% acetonitrile, 100 mM ammonium bicarbonate, 5 mM DTT, 16 h, 4 °C). Samples were then alkylated (40 mM chloroacetamide, 160 mM ammonium bicarbonate, 10 mM TCEP, 20 min, 70 °C), dehydrated in 100% acetonitrile, and air-dried at 37 °C, followed by digestion with trypsin (Promega). Digests were loaded onto Evotips and trypic peptides were eluted using the ‘30SD’ gradient via an Evosep One HPLC fitted with a 15 cm C18 column into a Lumos Tribrid Orbitrap Mass spectrometer via a nanospray emitter operated at 2200 V. The Orbitrap was operated in ‘Data Dependent Acquisition’ mode with precursor ion spectra acquired at 120k resolution in the Orbitrap, and MS/MS spectra in the ion trap at 32% HCD collision energy in ‘TopS’ mode. Dynamic exclusion was set to +/-10 ppm over 15 s, automatic gain control to ‘standard’ and max injection time to ‘Dynamic’. The vendors universal method was adopted to schedule the ion trap accumulation times.

Raw files were processed using MaxQuant 2.0.3.0 and Perseus with a recent download of the UniProt Homo Sapiens reference proteome (UP000005640), or *Escherichia coli* reference proteome (UP000000625), and the sequence of the FL-RNC based on UniProt ID Q27758_PHOPY, including N-terminal tag, and C-terminal linker and XBP1u+ or SecM stalling sequence. A decoy dataset of reverse sequences was used to filter false positives, with both peptide and protein FDR set to 1 %. Quantification of individual proteins was achieved using iBAQ (intensity based absolute quantification) and values were normalised to the mean iBAQ all 50S, or 70S proteins in each sample. Therefore, each protein can be quantified as a percentage per ribosome in the sample.

### Hydrogen/deuterium exchange-mass spectrometry

#### RNC labelling, quench and digestion

RNC labelling, quench and offline digest was carried out as previously described^19,65^ with some differences. 60 μL of 0.2-0.5 μM RNC, FLuc or ribosome was labelled in 20 mM HEPES-pH 7.5, 100 mM KCl, 5 mM Mg(OAc)_2_, 1 mM DTT and 0.2 M Arginine-HCl made up with water or D_2_O. Deuterated buffers contained 98% D_2_O. Labelling was initiated using 7K 0.5 mL Zeba spin desalting columns, pre-equilibrated with labelling buffer as per manufacturer’s instructions. Exchanged proteins were eluted by centrifugation at 1000g for 2 min at 25 °C, and incubated at 25 °C with 350 rpm shaking in an Eppendorf ThermoMixer. 50 μL of exchanged material was quenched at 3 min or 60 min with 20 μL of quench buffer (175 mM Na-phosphate, pH 2.1, 17.5 mM TCEP, 5 M GuHCl, adjusted to pH to 1.2 with orthophosphoric acid. The final pH after quench was 2.3) on ice, and immediately added to 20 μl of 50% pepsin POROS slurry^65^ pre-incubated at 10 °C. Offline RNC digest was carried out at 10 °C for 2 min with 450 rpm shaking, with brief vortexing every 30 s. The digested peptides were eluted through a 0.2 μm PVDF centrifugal filter (Durapore) at 0 °C for 30 s at 16,000g before flash freezing and storage at -80 °C, or immediate thaw and injection into a Synapt G2-Si (Waters).

Maximally-deuterated samples were prepared as previously described^85^. Undeuterated samples were digested as above, and hydrogen-containing solvent was removed overnight using a Savant ISS110 Speedvac (ThermoFisher). The pellets containing peptides were resuspended in 50 μL deuterated labelling buffer for 4 h at 40 °C, before adding 20 μL of quench buffer on ice, and incubating at 10 °C for 2 min before flash freezing in LN_2_.

Labelled and quenched peptides were thawed and immediately injected into a Waters HPLC coupled to Synapt G2-Si HDMS^E^ mass spectrometer. All chromatographic elements were held at 0 °C in the UPLC chamber. Peptides were trapped and desalted on a VanGuard pre-column trap (2.1 mm x 5 mm ACQUITY UPLC BEH C4 1.7 μm (Waters)) for 4 minutes at 200 μL.min^-1^. Peptides were eluted in a gradient of 3-35% acetonitrile in 0.1% formic acid over 25 minutes across at a flow rate of 90 μL.min^-1^, and separated using an ACQUITY UPLC HSS T3 1.8 um x 5 mm 1.8 μm LC column (Waters). Mass spectra were acquired using Waters Synapt G2-Si in ion-mobility (IMS) mode. The mass spectrometer was calibrated with direct infusion of a solution of Glu-fibrinopeptide at 50 fmol.μL^-1^ at 5 μL.min^-1^. A conventional electrospray source was used and the instrument was scanned over a 50 to 2000 m/z range. The instrument was configured as follows: capillary at 3 kV, trap collision energy at 4 V, sampling cone at 40 V, source temperature of 80 °C and desolvation temperature of 180 °C. All experiments were carried out under identical experimental conditions in triplicate and were not corrected for back exchange and therefore reported as relative.

After each injection, a trap column and LC wash was injected containing 1.5 M guanidinium-HCl, 4% (v/v) acetonitrile, 0.8% (v/v) formic acid, with peptide trapping time of 3 min and eluted with 3 repeat ramps of 5-95% ACN over 12 minutes at 90 μL.min^-1^.

#### Data analysis

Peptides were identified from replicate HDMS^E^ analyses of undeuterated control samples using PLGS 3.0.3 (Waters). Peptide masses were identified from searches using non-specific cleavage of a custom database containing the sequence of FL-RNC based on UniProt ID Q27758_PHOPY, and all ribosomal proteins identified and extracted from proteomics datasets. In addition, the database contained HSPA1A and EIF6 which were also identified in RNC samples. Searches used the following parameters: no missed cleavages, no PTMs (although methionine oxidation was allowed), a low energy threshold of 135, an elevated energy threshold of 35 and a lock mass window of 0.25 Da, 3 minimum fragment ion matches per peptide, 7 minimum fragment ion matches per protein. No false discovery rate correction was performed. Peptides identified in PLGS for each RNC were aggregated and further filtered using DynamX 3.0 (Waters) using a maximum sequence length of 40, 0.05 minimum products per amino acid and a minimum of 1 consecutive product identified. Each mass spectrum was manually inspected for spectral overlap and signal to noise, and peptides of unacceptable quality were removed. In addition, peptides were removed from NC analyses if they were assigned in control analyses of empty ribosomes. The relative amount of deuterium was determined with the software by subtracting the centroid mass of the undeuterated from the deuterated spectra at each timepoint for each condition. Spectra showing multimodal uptake kinetics were further analysed using HX-Express3^86^. Uptake values were used to generate all uptake graphs and difference maps using Graphpad Prism 10.1.1. See also Data S3, S5, S6 and S7.

## Data availability

Proteomic analysis used the Uniprot H. sapiens (UP000005640) or E. coli (UP000000625) reference proteome. All mass spectrometry data have been deposited to the ProteomeXchange Consortium via the PRIDE partner repository with the following dataset identifiers:

hRNC composition: PXD055069

XL-MS: PXD055251

HDX-MS analysis of hRNCs: PXD055280

bRNC composition: PXD055060

HDX-MS analysis of bRNCs: PXD055476

The cryoEM map has been deposited in the Electron Microscopy Data Bank (EMDB), and the coordinates have been deposited in the PDB under the following accession numbers: EMD-XXXX and PDB: XXXX

---

**Fig. S1.**
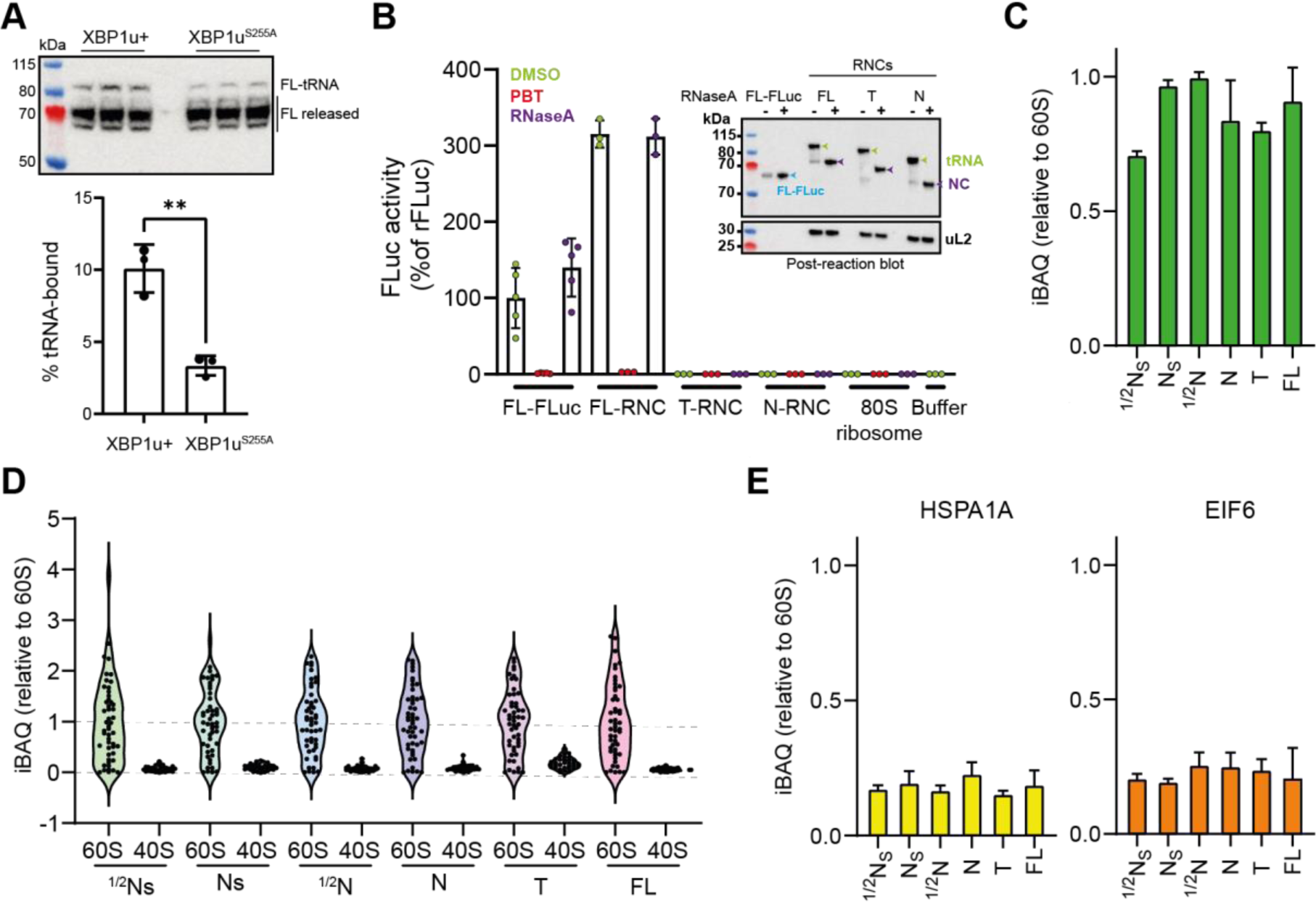
Quality control of FLuc RNCs. **A,** Comparison of stalling efficiency between XBP1u+ and XBP1u^S255A^. FL-RNC constructs prepared with either stalling sequence were expressed in Expi293 cells. Clarified lysates were resolved by SDS-PAGE and immunoblotted using an antibody against the N-terminus of FLuc. The intensity of the NC-tRNA band was measured using ImageJ, and expressed as a percentage of the total FLuc signal. Error bars, s.d.; n = 3 independent experiments. p= 0.0030**, two-tailed unpaired t-test. **B**, FL-RNC is enzymatically active. Luminescence activity is expressed as a percentage of the activity of FL-FLuc. Where indicated, samples were treated with RNase/EDTA to release the NC from ribosomes, or PBT to inhibit FLuc activity. Inset: control immunoblot (anti-FLuc) of post-reaction samples, showing that the NC-tRNA is intact. **C**, Fractional occupancy of FLuc in RNCs. Intensity-based absolute quantification (iBAQ) of NC peptides in each RNC, normalised to the average iBAQ of 60S ribosomal proteins. **D**, Quantification of ribosomal proteins in RNCs. iBAQ values for each protein were normalised to the average iBAQ of 60S ribosomal proteins. 60S and 40S proteins are grouped separately. **E**, Fractional occupancy of HspA1A (Hsp70) and EIF6 in RNCs. iBAQ values for HspA1A or EIF6 peptides in each RNC were normalised to the average iBAQ of 60S ribosomal proteins. See also Data S1.

**Fig. S2.**
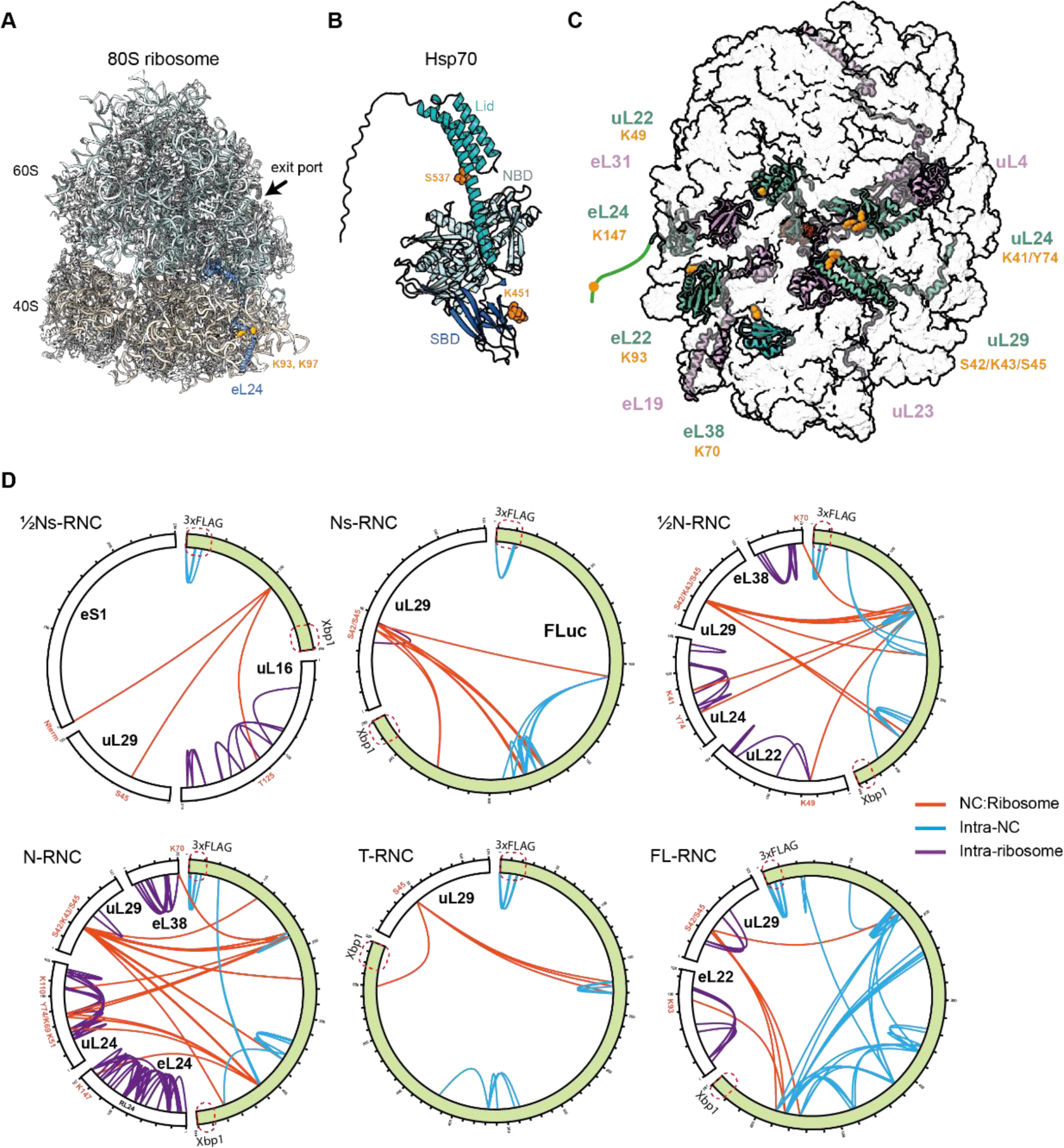
Crosslinking-mass spectrometry (XL-MS) analysis of RNCs. **A,** Hsp70 crosslink sites on the ribosome (PDB 8QOI^64^). Residues on eL24 (blue) which crosslink to Hsp70 are shown as orange spheres. The ribosomal exit port is indicated. **B**, Resides on Hsp70 that crosslinked to eL24 are shown as orange spheres. Hsp70 is shown in the “open” (domain-docked) conformation predicted by Alphafold2 (AF-P0DMV8-F1). **C**, NC crosslink sites on the ribosome. Ribosomal protein residues that crosslinked to any FLuc NC are shown as orange spheres on the structure of Ns-RNC. The N-terminal tail of eL24 containing K147 is not resolved in the structure. **D**, Map of RNC crosslinks identified by MS. NC:ribosome crosslinks (orange), intra-NC crosslinks (blue) and intra-ribosome crosslinks (purple) are shown. See also Data S2.

**Fig. S3.**
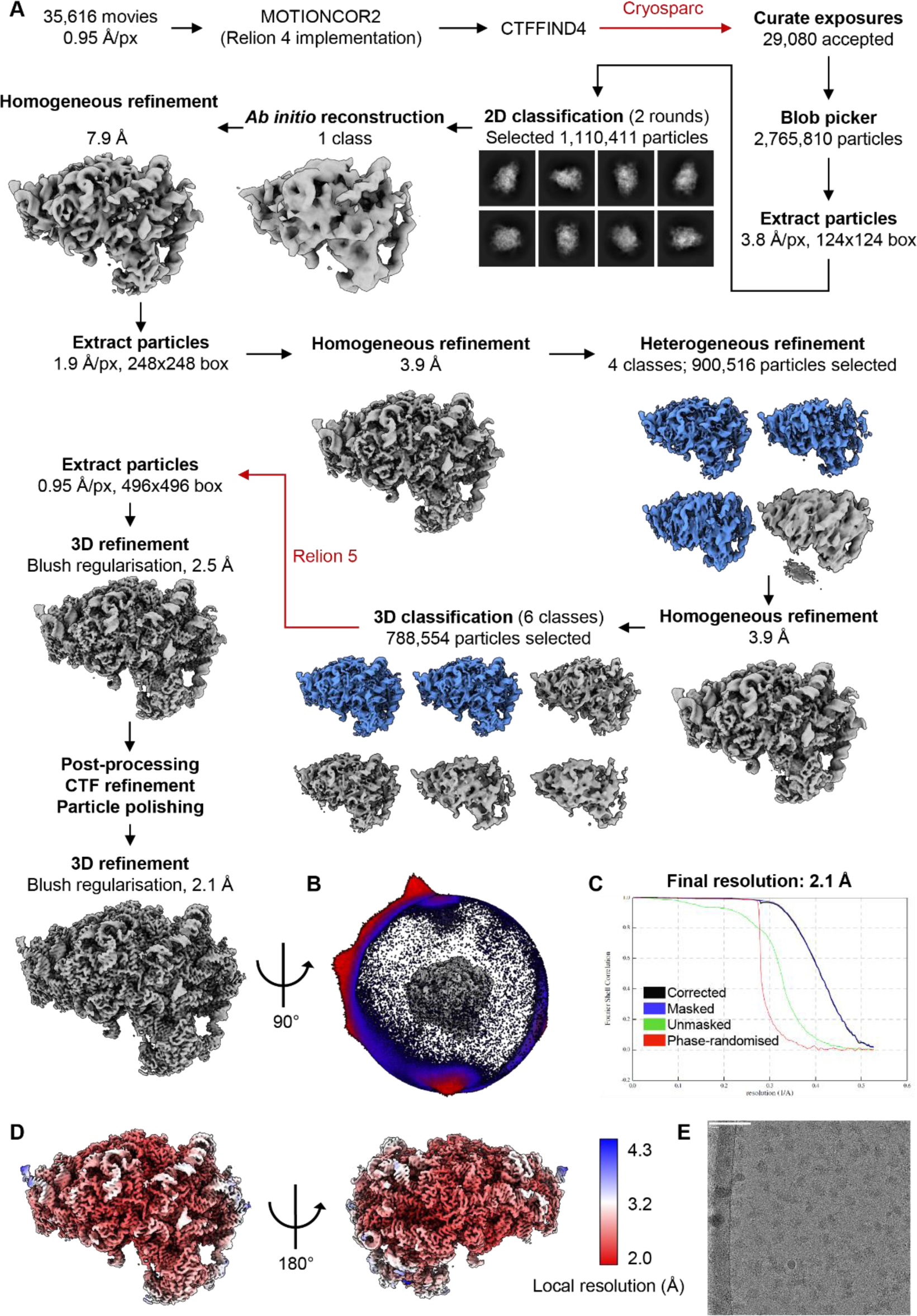
CryoEM image processing. **A,** Schematic of the cryoEM processing pipeline including representative 2D classes and the final reconstruction. **B,** Angular distribution of the final cryoEM map. **C,** Fourier Shell Correlation curve of the cryoEM reconstruction. **D,** CryoEM map coloured according to local resolution, from 2.0 Å (red) to 4.3 Å (blue). **E,** Representative micrograph (white scale: 75 nm).

**Fig. S4.**
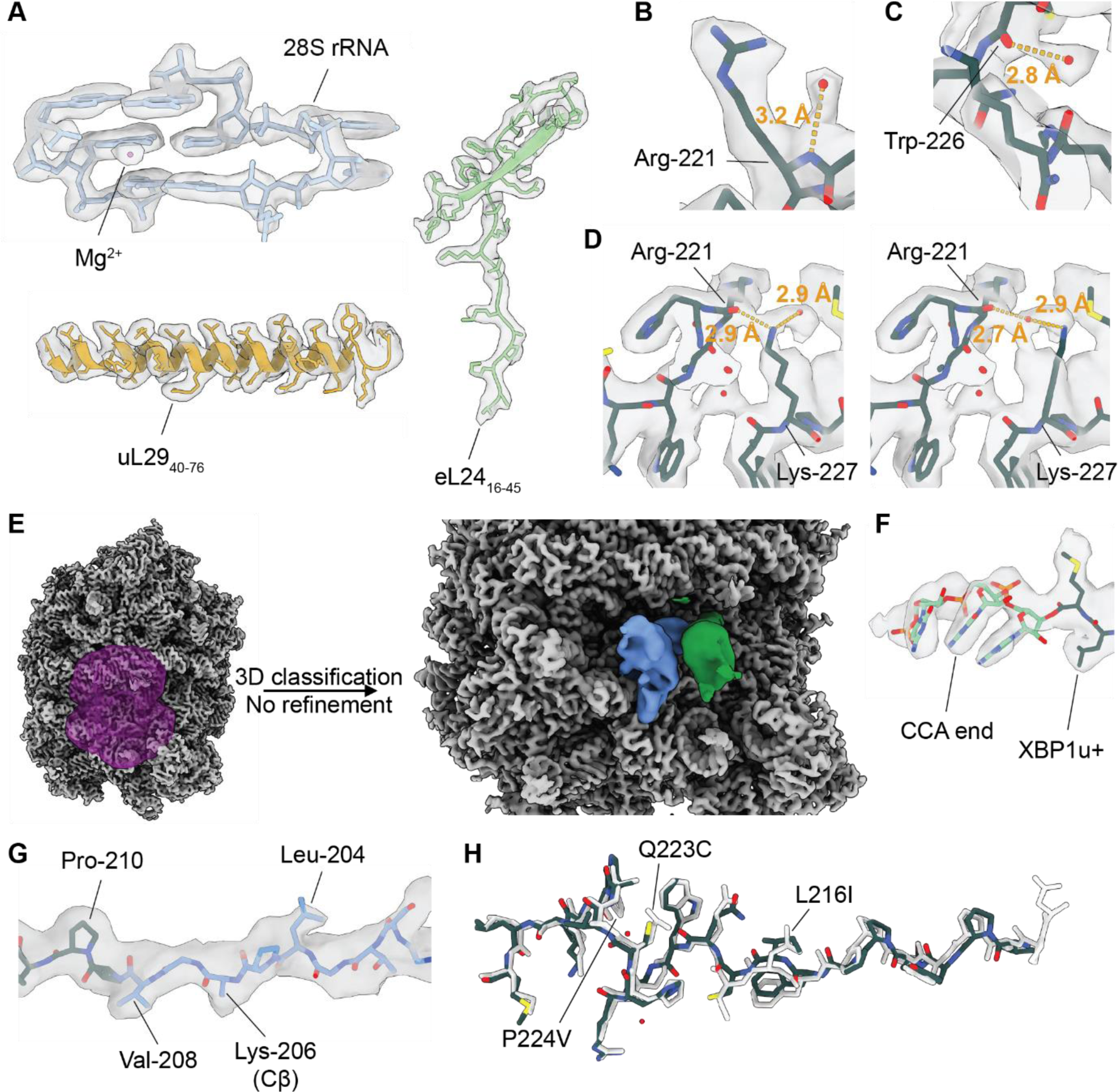
Quality of the cryoEM reconstruction and model building. **A,** Map-model overlay of example regions of the structure. **B,** Map-model overlay of XBP1u+ showing an ordered water molecule interacting with the backbone nitrogen of Arg-221. **C,** Map-model overlay of XBP1u+ showing an ordered water molecule interacting with the carbonyl oxygen of Trp-226. **D,** Map-model overlay of XBP1u+ with the side chain of Lys-227 modelled to form a hydrogen bond with the carbonyl oxygen of Arg-221 and an ordered water molecule (left) and with the ordered water molecule placed between Arg-221 and Lys-227 (right). As the position of the water molecule could not be unambiguously determined, it was not modelled in the final structure. Distances in orange in B, C and D correspond to the hydrogen bonds marked by the orange lines. **E,** 3D classification with a mask around the tRNA sites (purple) shows P-tRNA flexibility. The P-tRNA density from two example classes (blue, green) is shown in the context of the consensus map (grey). **F,** Continuous density between the P-tRNA CCA end and XBP1u+ in the consensus map. **G,** Map-model overlay of the nascent chain with the side chain densities used to trace the backbone labelled. Due to the lack of density, the Lys-206 was trimmed to its Cβ carbon. **H,** Structural alignment of the XBP1u^S225A^ arrest peptide (white) and the stalling-enhanced XBP1u+ (coloured).

**Fig. S5.**
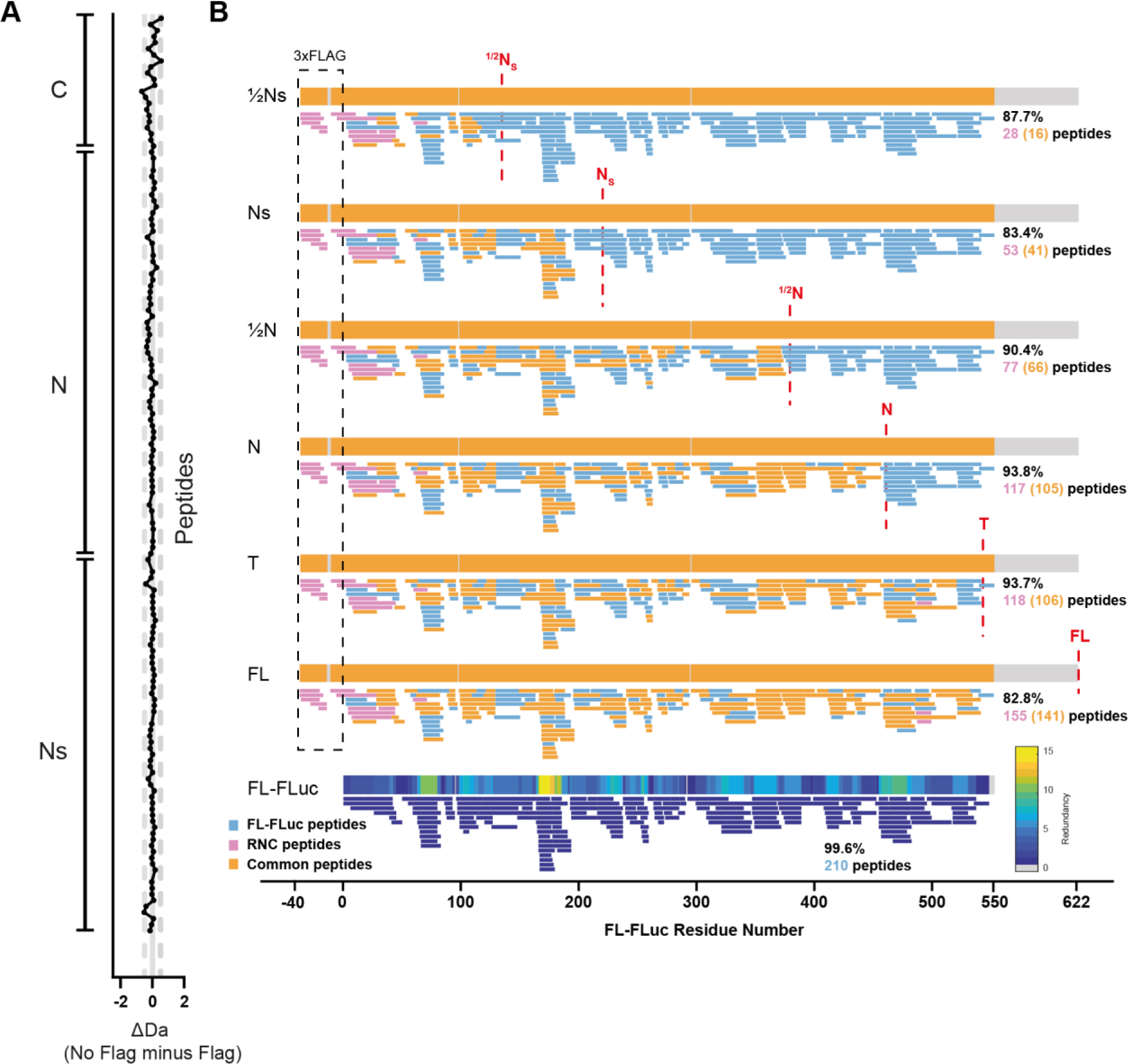
HDX-MS analysis of RNCs. **A,** FLAG tag does not measurably affect the conformation of FL-RNC. FL-RNC was treated with 3C protease to remove the 3xFLAG tag, then analysed by HDX-MS using a 3 min deuteration pulse. Grey dashed lines indicate ±0.5 Da. **B**, Peptide coverage maps for RNCs and FL-FLuc. Peptides found in FL-FLuc are coloured blue, peptides found in RNCs are coloured pink, and peptides common to both an RNC and FL-Fluc are coloured orange.

**Fig. S6.**
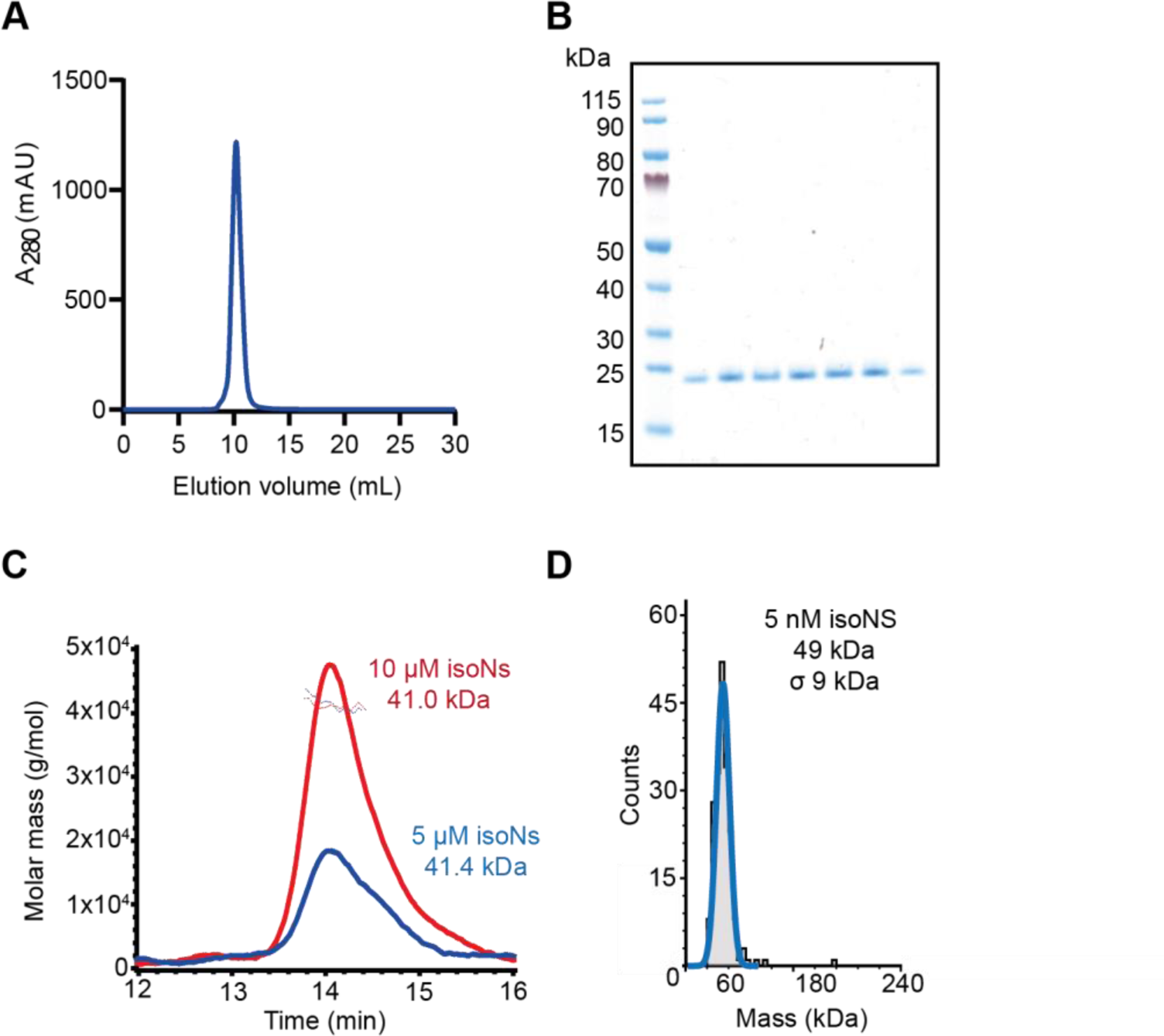
Characterisation of isolated Ns (isoNs). **A,** Gel filtration analysis using a Superdex 75 column indicates that purified isoNs is homogeneous. **B,** Fractions from the gel filtration peak in **A** were resolved by SDS-PAGE and stained with Coomassie. **C**, SEC-MALLS analysis of isoNs at 0.1 or 0.2 mg/mL (∼5 or 10 µM) indicates that isoNs is dimeric. Chromatograms are differential refractometer output, and points represent calculated molar mass. The expected mass of the monomer is ∼ 21.5 kDa. **D**, Analysis of 5 nM isoNs by mass photometry indicated an absolute mass of 49 kDa, σ 9 kDa, consistent with a dimer. Note that a monomeric peak at ∼21 kDa is not expected to be resolved.

**Fig. S7.**
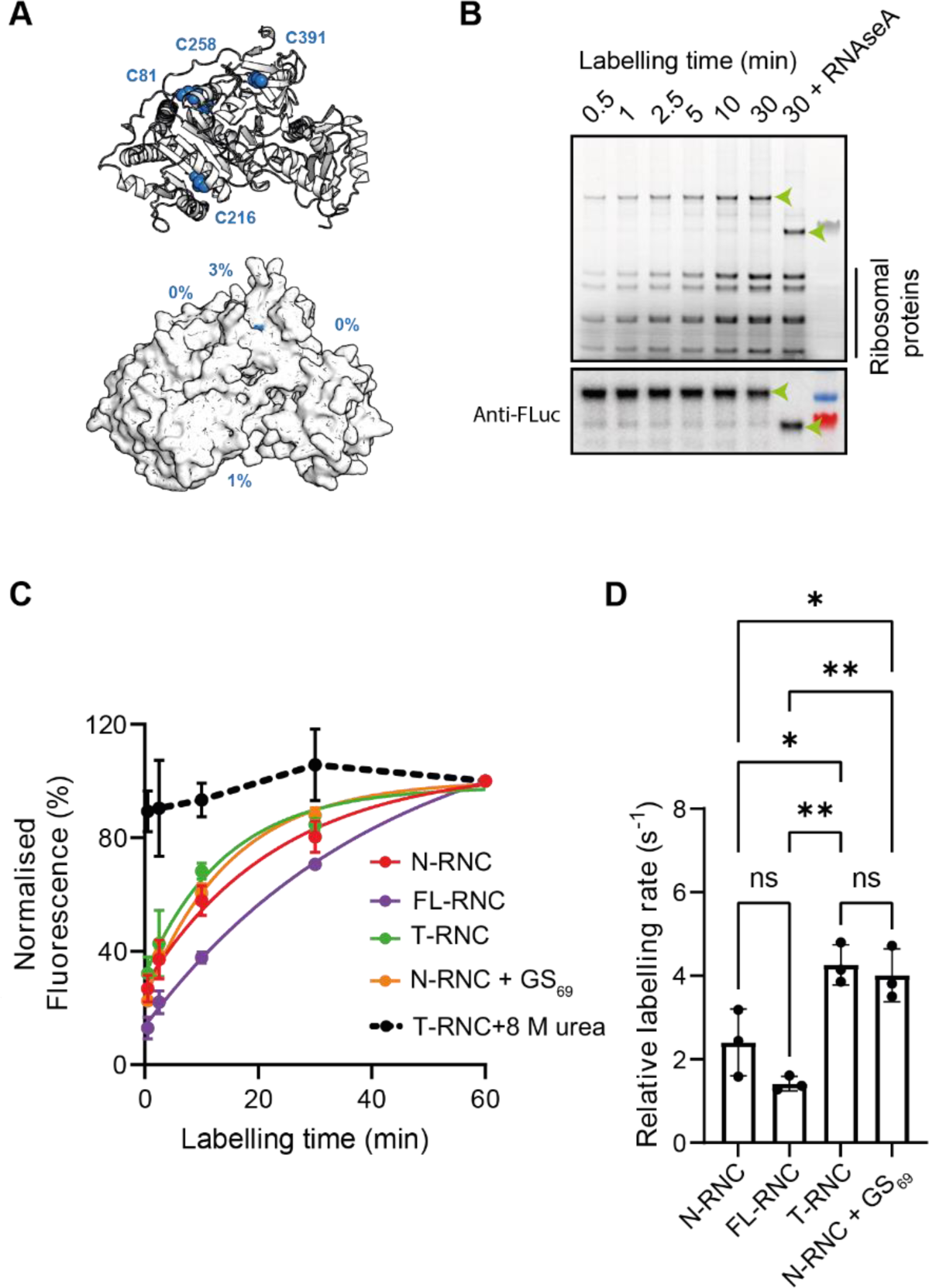
Analysis of NC conformation by cysteine labelling. **A,** Cys are buried in the native state of FL-FLuc. Cys residues in FLuc are shown as blue spheres. % solvent accessibility of each residue was calculated using PyMol. **B**, Fluorescein-5-malemide (F5M) labelling of T-RNC. Reactions were resolved by SDS-PAGE and imaged for in-gel fluorescence. The band corresponding to the NC is indicated using a green arrow. Below, the same gel was probed using an antibody against FLuc as a loading control. **C**, Cys labelling kinetics. The degree of NC labelling was quantified from gels as in **B** using ImageJ, and normalised to the maximum value. As a control, T-RNC was denatured in 8 M urea before labelling. Data were fit to a single-exponential equation and the resulting labelling rates are shown in **D**. Error bars, s.d.; n = 3 independent experiments. P-values were determined using a 1-way ANOVA. N-RNC vs T-RNC, p= 0.0167*; N-RNC vs FL-RNC, p= 0.2264 ns; N-RNC vs N-RNC+GS69, p= 0.0351*; T-RNC vs FL-RNC, p=0.0013**; T-RNC vs N-RNC+GS69, p= 0.9461 ns; FL-RNC vs N-RNC+GS69, p= 0.0023**.

**Fig. S8.**
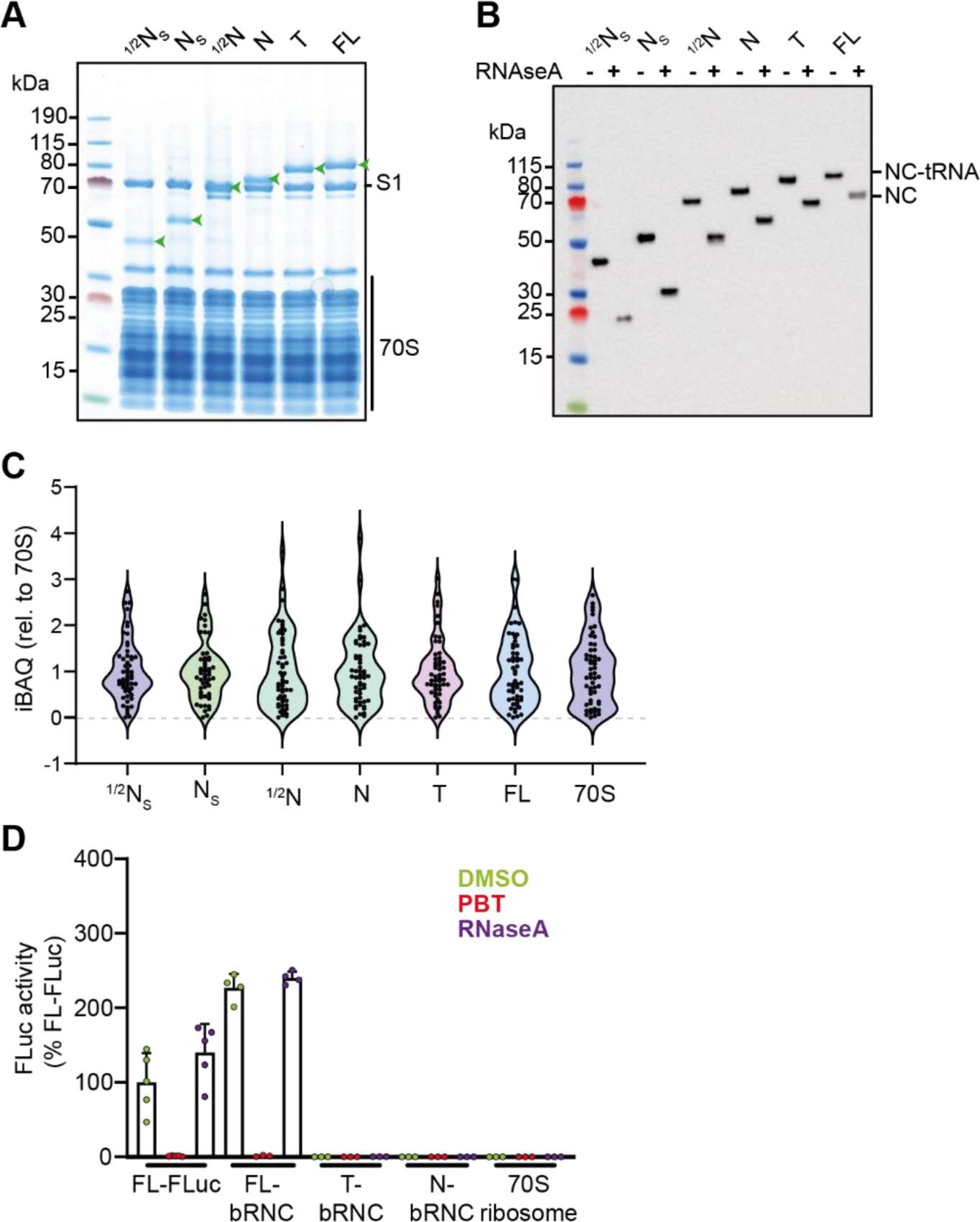
Quality control of FLuc bacterial RNCs (bRNCs). **A,** Coomassie-stained SDS-PAGE of purified bRNCs. Bands corresponding to 70S ribosomal proteins, ribosomal protein S1, and the NC linked to peptidyl-tRNA (green arrows) are indicated. **B**, Anti-FLuc immunoblot of purified RNCs. RNase/EDTA treatment confirms that the NCs are covalently linked to peptidyl-tRNA. **C**, Spread of iBAQ values for 70S ribosomal proteins in each bRNC. Purified empty ribosomes (70S) are shown as a control. **D**, FL-bRNC is enzymatically active. Luminescence activity is expressed as a percentage of the activity of FL-FLuc. Where indicated, samples were treated with RNaseA/EDTA to release the NC from ribosomes, or PBT to inhibit FLuc activity. See also Data S4.

**Fig. S9.**
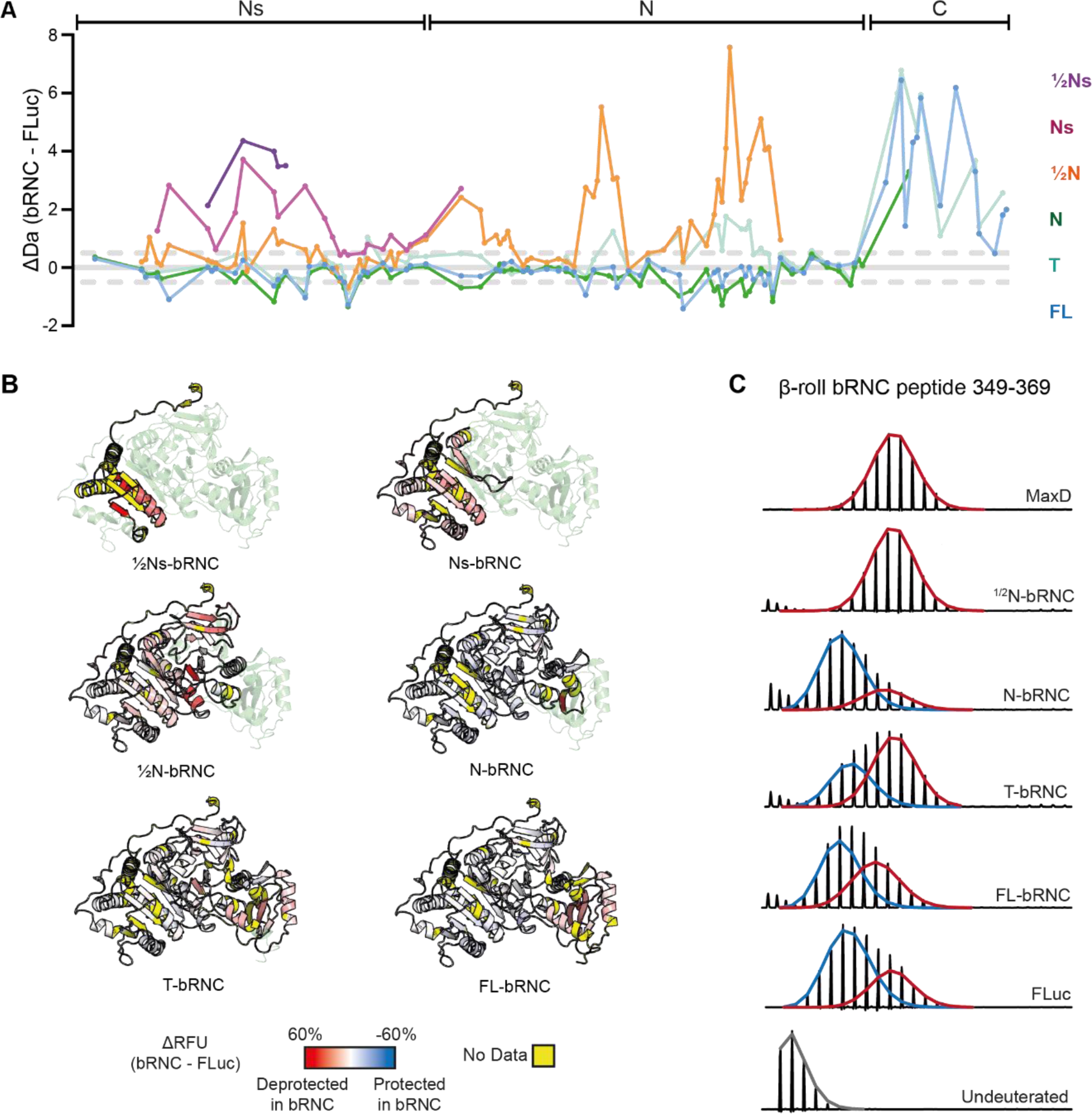
HDX-MS analysis of bRNCs. **A,** Difference in deuterium uptake, after 3 min deuteration, between FLuc NCs on the bacterial ribosome and native FL-FLuc. Larger values indicate more deuteration of NCs compared to FL-FLuc. Grey dashed lines indicate ±0.5 Da. Data represent the mean of at least 3 independent experiments. **B**, Difference in relative fractional uptake (ΔRFU), after 3 min deuteration, between each NC and FL-FLuc. Data are mapped onto the Alphafold2 model FL-FLuc. Darker red indicates increased deuteration of NCs compared to FL-FLuc. Regions without peptide coverage are coloured yellow. **C**, Mass spectral envelopes for peptide 349-369 in the β-roll of FLuc and bRNCs. Isotopic distributions were fit to single or bimodal gaussian distributions using HX-Express3.0 ^62^. The high-exchanging population (deprotected) is coloured red and the low-exchanging population (protected) is coloured blue. Except for the maximally deuterated sample (MaxD), samples were deuterated for 3 min. See also Data S5.

**Fig. S10.**
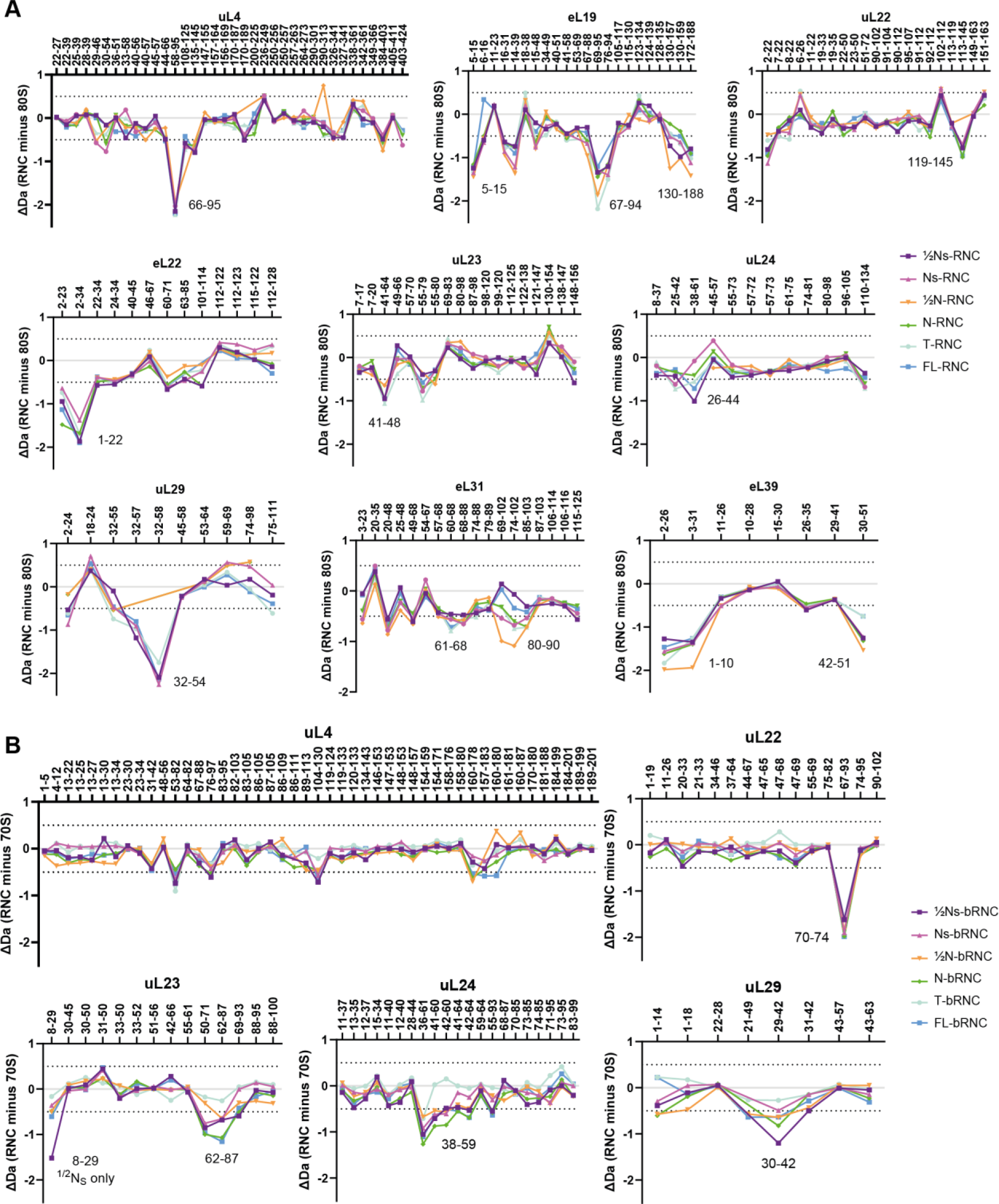
HDX-MS analysis of human and bacterial ribosomal proteins. **A,** Difference in deuterium uptake, after 3 min deuteration, between ribosomal proteins in empty human 80S ribosomes and different human RNCs. Dashed lines indicate ±0.5 Da. **B,** As in **A**, for bRNCs compared to empty 70S ribosomes. See also Data S6 and Data S7.

